# *Orientia tsutsugamushi* Ank5 directs ubiquitination and proteasomal degradation of NLRC5 to inhibit major histocompatibility complex class I expression

**DOI:** 10.1101/2023.12.04.569938

**Authors:** Haley E. Adcox, Jason R. Hunt, Kyle G. Rodino, Andrew K. Ottens, Jason A. Carlyon

## Abstract

How intracellular bacteria subvert the major histocompatibility complex (MHC) class I pathway is poorly understood. Here, we show that the obligate intracellular bacterium *Orientia tsutsugamushi* uses its effector protein, Ank5, to orchestrate proteasomal degradation of the MHC class I gene transactivator, NLRC5. Ank5 uses a tyrosine in its fourth ankyrin repeat to bind the NLRC5 N-terminus while its F-box directs host SCF complex ubiquitination of K1194 in the leucine-rich repeat region that dictates NLRC5 susceptibility to *Orientia*- and Ank5-mediated degradation. The ability of *O. tsutsugamushi* strains to degrade NLRC5 correlates with *ank5* genomic carriage. Ectopically expressed Ank5 that can bind but not degrade NLRC5 protects the transactivator during *Orientia* infection. Thus, Ank5 is an immunoevasin that uses its bipartite architecture to rid host cells of NLRC5 and MHC class I molecules. This study offers insight into how intracellular pathogens can impair MHC class I expression to benefit their survival.

## INTRODUCTION

Major histocompatibility complex (MHC) class I antigen presentation is essential for adaptive immunity against intracellular microbes.^1^ Yet many intracellular pathogens have evolved strategies to undermine this pathway. Such approaches have been well studied for viruses, but considerably less is known about how intracellular bacteria subvert MHC class I-mediated immunity.^2-7^ *Orientia tsutsugamushi* is an obligate intracytosolic bacterium and symbiont of trombiculid mites.^8^ It causes scrub typhus, a disease known for centuries to be endemic to the Asia-Pacific where an estimated one million cases occur annually.^8,9^ Locally acquired cases in Africa, the Middle East, and South America and detection of *Orientia* species in mites in the United States signify scrub typhus as a globally emerging health threat.^10-17^ *O. tsutsugamushi* infects leukocytes at the mite feeding site and disseminates to invade endothelial cells of multiple organs.^18,19^ Acute scrub typhus symptoms are non-specific and include fever, rash, headache, and lymphadenopathy.^20,21^ If untreated, *O. tsutsugamushi* infection of endothelial cells can progress to systemic vascular collapse, organ failure, and death.^22,23^ *O. tsutsugamushi* strains are extensively diverse in terms of virulence in humans and animal models.^24-31^ The molecular bases for variations in the virulence capacities of different *O. tsutsugamushi* strains are poorly defined.

MHC class I molecules and CD8^+^ T cells are important for restraining *O. tsutsugamushi* growth and protecting against lethal infection in mice.^25,32,33^ Moreover, CD8^+^ T cell numbers are elevated during the convalescent but not acute phase in scrub typhus patients.^34^ Hence, *O. tsutsugamushi* must counter the MHC class I pathway to maximize its survival in animal and human hosts. NLRC5 (NOD-, LRR- and CARD-containing 5/class I transactivator) is a master regulator of MHC class I gene expression.^35,36^ It is upregulated by interferon-γ (IFNγ) and is the sole transactivator of MHC class I gene expression in endothelial cells and other non-professional antigen presenting cells (APCs).^36-40^ The *O. tsutsugamushi* Ikeda strain lowers NLRC5 levels in non-professional APCs, even in the presence of IFNγ, to inhibit MHC class I gene expression and reduce the amounts of MHC class I molecules on the cell surface.^41^ Ikeda, isolated from a patient in Japan, causes severe disease in humans and is fatal in laboratory mice.^42-47^ It is the only microbe known to modulate MHC I cellular levels by targeting NLRC5, although the mechanism is uncharacterized.

Ikeda and all other sequenced *O. tsutsugamushi* strains encode among the largest microbial arsenals of ankyrin repeat (AR)-containing effector proteins (Anks).^42,48-52^ ARs are protein-protein interaction motifs that are conserved across the Tree of Life.^51^ Their acquisition by intracellular microbes is linked to increased virulence.^53,54^ Most *O. tsutsugamushi* Anks carry a C-terminal eukaryotic-like F-box domain, which interacts with host Skp1 (S-phase kinase-associated protein 1) and Cul1 (cullin 1) to nucleate the SCF (Skp1-Cul1-F-box) E3 ubiquitin ligase complex that mediates polyubiquitination of substrates to destine them for 26S proteasomal degradation.^55,56^ This AR-F-box bipartite architecture is exhibited by numerous other intracellular bacterial effectors and viral proteins.^53,57-62^ Strikingly, however, nearly all of the host cell target proteins that get ubiquitinated and degraded due to interacting with microbial F-box-containing Anks remain unidentified. Thus, understanding of this evolutionarily conserved intracellular microbial pathogenesis strategy is limited.

Here, we report that *O. tsutsugamushi* Ikeda Ank5 mediates NLRC5 degradation to reduce MHC class I cellular levels. Specifically, the fourth AR of Ank5 binds the NLRC5 N-terminus, enabling the effector’s F-box to direct SCF-mediated ubiquitination of a lysine in the NLRC5 leucine-rich repeat (LRR) domain. Ectopic overexpression of a recombinant version of Ank5 capable of binding but not degrading NLRC5 protects the transactivator from degradation during *Orientia* infection, further establishing the respective contributions of the AR and F-box domains to Ank5 function. *O. tsutsugamushi* UT76, another strain that carries *ank5*, is capable of lowering NLRC5 and MHC class I levels, whereas the Karp strain, which lacks *ank5*, cannot. Our findings elucidate a detailed mechanism of action for the first identified microbial immunoevasin that subverts the MHC class I pathway by degrading NLRC5.

## RESULTS

### *O. tsutsugamushi* promotes NLRC5 degradation by the 26S proteasome

*O. tsutsugamushi* Ikeda (hereafter referred to as *O. tsutsugamushi* unless discussed in the context of other strains) decreases NLRC5 protein levels in non-professional APCs even though it induces *nlrc5* transcription,^41^ which suggests that the pathogen targets the transactivator at a posttranscriptional step. As further support for this hypothesis, *O. tsutsugamushi* reduces levels of IFNγ-induced endogenous NLRC5 as well as ectopically expressed Flag- and myc-tagged NLRC5 (Figure 1A-1D). HeLa cells were used for this and other experiments herein because they are non-professional APCs that constitutively express MHC class I genes in an exclusively NLRC5-dependent manner.^36,63^ HeLa cells are also established models for studying *O. tsutsugamushi*-host interactions, and their amenability to transfection allows for assessing if ectopically expressed oriential virulence factors phenocopy aspects of infection.^41,64-73^

**Figure 1.**
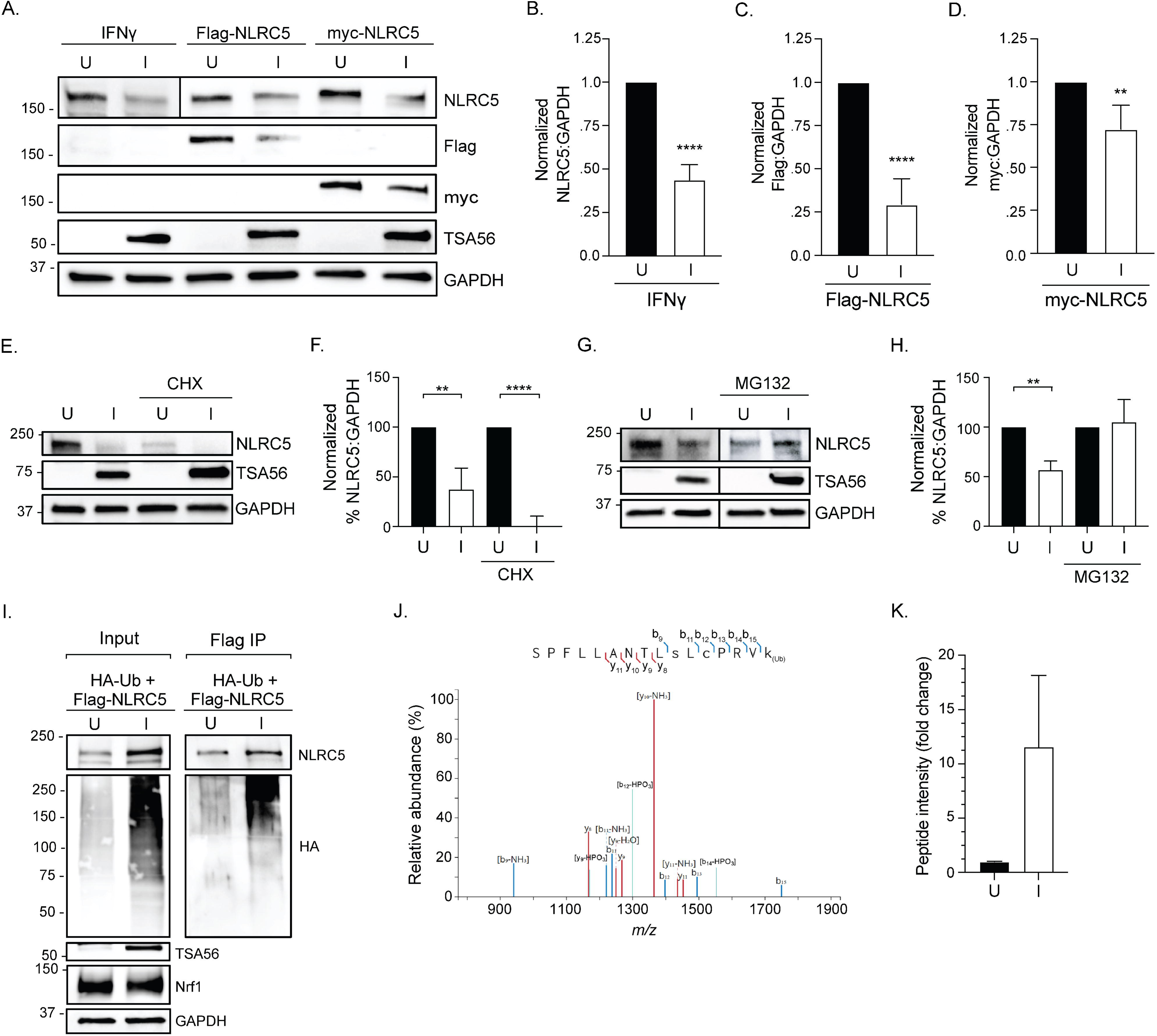
*O. tsutsugamushi* promotes ubiquitination and proteasomal degradation of NLRC5. (A–D) HeLa cells were mock transfected and treated with IFNγ or transfected to express Flag-NLRC5 or myc-NLRC5 and then mock infected [U] or infected with *O. tsutsugamushi* [I]. (A) Whole cell lysates were analyzed by immunoblotting. Vertical lines between bands indicate where the blot was cropped or imaged separately. (B–D) NLRC5 (B), Flag (C), or myc (D) densitometric signal was divided by the corresponding GAPDH signal and normalized to the U densitometric value per sample. Data are means ± SD (n = 4 to 5R). An unpaired *t*-test determined statistical significance among pairs. (E–H) HeLa cells were mock infected [U] or infected with *O. tsutsugamushi* [I] and treated with either cycloheximide (CHX) (E and F) or MG132 (G and H) for 16 hr prior to collection. (E and G) Whole cell lysates were analyzed by immunoblotting. Vertical lines between bands indicate where the blot was cropped or imaged separately. (F and H) NLRC5 densitometric values were divided by the corresponding GAPDH densitometric signal. The quotients were normalized to U densitometric values and multiplied by 100. Data are means ± SD (n = 3). An unpaired *t*-test determined statistical significance among pairs. (I) HeLa cells were transfected to co-express Flag-NLRC5 and HA-Ub, mock infected [U] or infected with *O. tsutsugamushi* [I] and treated with MG132 for 24 hr prior to collection. Input lysates and immunoprecipitated Flag-NLRC5 (Flag IP) complexes were analyzed by immunoblotting (n = 3). (J and K) HeLa cells were transfected to express Flag-NLRC5, mock infected [U] or infected with *O. tsutsugamushi* [I] and treated with MG132 for 24 hr prior to collection. Flag-NLRC5 was immunoprecipitated from collected lysates, eluted, and the eluted complex was analyzed by mass spectrometry. (J) Fragmentation spectra of the post-translationally modified _1179_SPFLLANTLSLCPRVK_1194_ peptide. Spectra-matched b- (blue) and y-ion (red) fragments are displayed and summarized by the trypsinized peptide sequence inset above. (K) Peptide counts presented as mean fold change ± SEM (n = 3). **, p < 0.01; ****, p < 0.0001.

To determine if *O. tsutsugamushi* prevents translation of nascent NLRC5 protein or targets NLRC5 for proteasomal degradation, HeLa cells were infected or not followed by the addition of cycloheximide, a eukaryotic protein synthesis inhibitor, or MG132, a 26S proteasome inhibitor. At 24 hr post-infection (p.i.), NLRC5 levels were decreased by approximately 50% in vehicle treated cells (Figure 1E–1H). *O. tsutsugamushi*-mediated NLRC5 reduction persisted in cycloheximide-treated cells to the point that NLRC5 was undetectable (Figure 1E and 1F). In contrast, NLRC5 levels were equivalent in MG132 treated infected and uninfected cells. Thus, *O. tsutsugamushi* promotes NLRC5 degradation by the 26S proteasome (Figure 1G and 1H).

### NLRC5 is ubiquitinated at K1194 in *O. tsutsugamushi* infected cells

Proteins destined for proteasomal degradation are marked by the covalent addition of polyubiquitin. To assess whether *O. tsutsugamushi* promotes NLRC5 ubiquitination, HeLa cells transfected to co-express hemagglutinin (HA)-ubiquitin (Ub) and Flag-NLRC5 were infected for 24 hr, after which MG132 was added for an additional 24 hr to enrich for ubiquitinated proteins. Lysates were collected followed by immunoprecipitation of Flag-NLRC5. Successful MG132 treatment was confirmed by the upregulation of Nrf1^74^ (Figure 1I). NLRC5 levels were markedly increased in MG132 treated infected cells, likely due to the combinatorial effect of upregulated *nlrc5* expression in response to *O. tsutsugamushi* infection,^41^ Flag-NLRC5 overexpression, and 26S proteasome inhibition. Indeed, levels of HA-ubiquitinated proteins overall and HA-ubiquitinated NLRC5 were more abundant in MG132 treated infected cells versus MG132 treated uninfected cells (Figure 1I).

To identify the NLRC5 residue(s) that becomes ubiquitinated during *O. tsutsugamushi* infection, HeLa cells expressing Flag-NLRC5 were infected and treated with MG132. Input lysates (Extended Data Figure 1) were subjected to immunoprecipitation to recover Flag-NLRC5 and interacting proteins. Eluted complexes were processed for liquid chromatography with tandem mass spectrometry (LC-MS/MS) and peptide fragments were examined for diglycine motifs, the post-tryptic mark of ubiquitination. Sequences were mass matched ± four parts per million and aligned for analysis. The proteome precipitated by Flag-NLRC5 from *O. tsutsugamushi* infected cells was three times more complex than that from uninfected cells (Extended Data Figure 2A). STRING (Search Tool for the Retrieval of Interacting Genes/Proteins)^75^ analysis of the infection specific NLRC5 interacting proteome revealed an enrichment for 26S proteasome subunits, proteins involved in unfolded protein binding, and other gene ontology categories (Extended Data Table 1). Filtering the infection-specific interactome to display only known interactions revealed that none of these proteins have been reported to form interacting networks with NLRC5 (Extended Data Figure 2B). Of note, recovery of 26S proteasome subunits and proteins involved in unfolded protein binding is consistent with NLRC5 becoming marked for proteasomal degradation. Analysis of NLRC5 tandem mass spectra revealed the peptide _1179_SPFLLANTLSLCPRVK_1194_ to have b- and y-ions specifically positioning a diglycine motif on residue K1194 (Figure 1J), which was recovered at 10-fold greater abundance from infected versus uninfected samples (Figure 1K). Altogether, these data demonstrate that *O. tsutsugamushi* promotes ubiquitination of NLRC5 on K1194, which leads to its degradation by the 26S proteasome.

### Ank5 is a putative NLRC5 binding partner that *O. tsutsugamushi* transcriptionally upregulates prior to extensive intracellular growth

NLRC5 degradation in *O. tsutsugamushi* infected cells is bacterial protein synthesis-dependent,^41^ suggesting that an oriential protein mediates this phenomenon. *O. tsutsugamushi* Ikeda Ank5 is a 345-amino acid effector protein of unknown function that has an N-terminal domain consisting of four tandemly arranged ARs and a C-terminal F-box that occurs within a PRANC (<u>p</u>ox proteins <u>r</u>epeats of <u>an</u>kyrin-<u>C</u> terminal) domain (Extended Data Figure 3A), the latter of which is a feature shared only with other *O. tsutsugamushi* Anks and Anks of vertebrate poxviruses, *Wolbachia* species, and wasps that *Wolbachia* species infect.^42,50,53,55,69,76,77^ The Ikeda chromosome encodes three identical Ank5 copies (OTT_RS01000, OTT_RS01950, and OTT_RS02325).^42,50^ We assessed *ank*5 transcription over the course of *O. tsutsugamushi* growth in HeLa cells and EA.hy926 human vascular endothelial cells. Bacterial replication in HeLa cells lagged for the first 24 hr after which it pronouncedly increased, whereas in EA.hy926 cells replication initiated during the first 12 hr and became logarithmic between 12 and 24 hr (Extended Data Figure 3B). Although *ank5* transcripts were detectable throughout infection of both cell types, significant *ank5* upregulation occurred at 8 and 12 hr p.i. in HeLa cells and at 2, 4, and 8 hr p.i. in EA.hy926 cells (Extended Data Figure 3C and 3D), indicating that *O. tsutsugamushi* strongly expresses *ank5* prior to its exponential growth phase.

A yeast two-hybrid screen was performed using Ank5 lacking residues 301-344 (Ank5ΔF-box) as bait so that known F-box dependent interacting partners Skp1 and Cul1^55^ would not be overrepresented among the prey proteins. One of the putative Ank5ΔF-box interactors was NLRC5 (Extended Data Table 2), a conspicuous finding given the effect of Ikeda on NLRC5 and because NLRC5 was not identified as a potential interactor of other Ikeda Anks by prior yeast two-hybrid screenings.^64,66^

### The ability of *O. tsutsugamushi* strains to reduce NLRC5 cellular levels correlates with genomic carriage of *ank5*

Because *O. tsutsugamushi* is genetically intractable^78^ we compared strains that do or do not carry *ank5* for the ability to lower host cell NLRC5 and MHC class I levels. Of the more than 30 antigenically distinct *O. tsutsugamushi* strains, nine have been fully sequenced and the *ank* genomic content has been determined for eight.^42,48,49,52,79^ Two of these strains, Ikeda and UT76 (NZ_LS398552.1), the latter of which originated from a patient in Thailand,^80^ carry *ank5*.^52^ A search performed using the National Center for Biotechnology Information (NCBI) Nucleotide Basic Local Alignment Search Tool (BLASTN) (https://blast.ncbi.nlm.nih.gov/Blast.cgi) with *ank5* (OTT_RS01000) unique nucleotides 400-546 as query identified an additional homolog in Wuj/2014 (NZ_CP044031.1), an isolate that had been recovered from a patient in China. The UT76 and Wuj/2014 *ank5* homologs are single-copy and exhibit 96–97% nucleotide and 90–92% amino acid identity with Ikeda Ank5 (Extended Data Table 3).

Classical MHC class I molecules consist of a heavy chain (HLA-A, -B, or -C) and β2-microglobulin (β2M).^81^ UT76 infected HeLa cells exhibited marked decreases in total NLRC5, HLA-ABC, and β2M along with cell surface MHC class I (Figure 2A to 2H), mirroring results reported for Ikeda infected cells.^41^ *O. tsutsugamushi* Karp, which does not encode Ank5,^52^ induced an approximate three-fold increase in NLRC5 levels and did not alter HLA-ABC and β2M amounts (Figure 2I to 2L). Yet, Karp infected cells still exhibited significantly reduced MHC class I surface levels (Figure 2M to 2P). Thus, *O. tsutsugamushi* Ikeda and UT76, which carry *ank5*, reduce NLRC5 levels during infection while Karp, which lacks *ank5*, does not.

**Figure 2.**
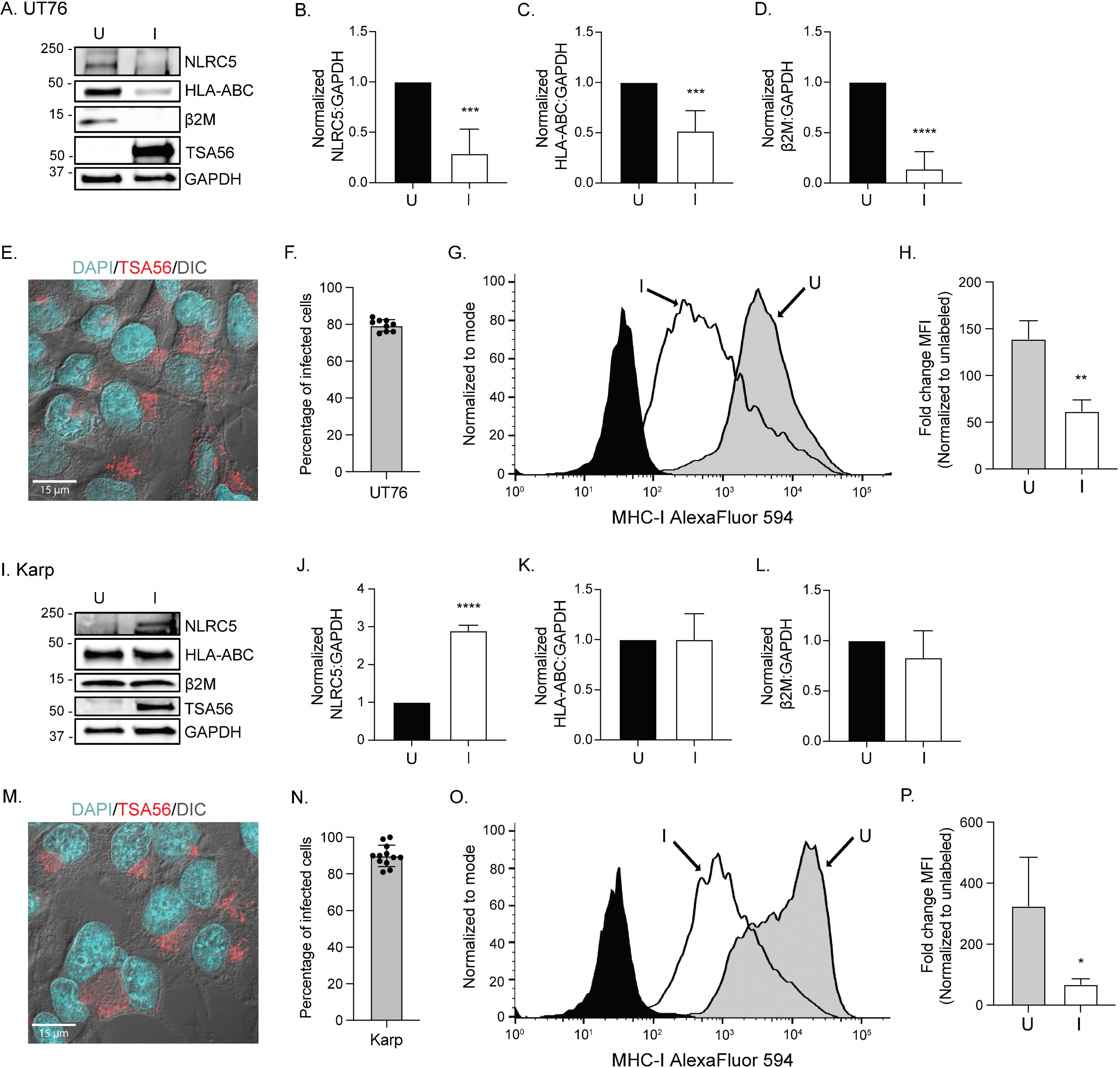
*O. tsutsugamushi* strain UT76, but not *O. tsutsugamushi* strain Karp, decreases cellular levels of NLRC5 and MHC class I components, yet both strains decrease surface MHC class I levels. (A–P) HeLa cells were mock infected [U] or infected with *O. tsutsugamushi* strain UT76 [I] (A– H) or *O. tsutsugamushi* strain Karp [I] (I–P) and treated with IFNγ. (A and I) Whole cell lysates were analyzed by immunoblotting. (B–D; J–I) The densitometric signal value of NLRC5 (B and J), HLA-ABC (C and K), or ϕ32M (D and I) was divided by the corresponding GAPDH densitometric signal value and normalized to the densitometric signal level of uninfected cells (black bars). Data are means ± SD (n = 3 to 5). An unpaired *t*-test determined statistical significance among pairs. (E and F; M and N) Percentage of infection was visualized by immunofluorescent microscopy (E and M) and quantified (F and N). (G and H; O and P) Surface levels of MHC class I from uninfected and infected cells were analyzed by flow cytometry. (G and O) Representative histograms of MHC class I levels normalized to mode. Unlabeled (black) samples served as the control. (H and P) The mean fluorescence intensity of each sample was divided by that of its unlabeled control per replicate. Data are means ± SD (n = 3 to 4). An unpaired *t*-test determined statistical significance among pairs. ***, p < 0.001; ****, p < 0.0001.

### Ank5 binds and decreases levels of NLRC5

To validate the predicted Ank5-NLRC5 interaction, HeLa cells were transfected to express Flag-tagged Ikeda Ank5 and treated with IFNγ. As a specificity control, cells were also transfected to express Flag-tagged Ank9, an *O. tsutsugamushi* effector that destabilizes the Golgi and is not predicted to bind NLRC5.^64^ Western blot analysis of input lysates and Flag antibody-immunoprecipitated complexes confirmed that NLRC5 was upregulated in IFNγ treated cells and precipitated by Flag-Ank5 but not Flag-Ank9 (Figure 3A). Conspicuously, NLRC5 levels were 64–70% lower in IFNγ-treated cells expressing Flag-Ank5 compared to mock transfected or Flag-Ank9 expressing cells (Figure 3A and 3B). Flag-Ank5 also significantly reduced NLRC5 levels in transfected EA.hy926 cells to phenocopy the effect observed in *O. tsutsugamushi* infected EA.hy926 cells (Extended Data Figure 4A–4D). NLRC5 was nearly depleted in infected EA.hy926 cells at 24 hr p.i. (Extended Data Figure 4C and 4D), which is 24 to 48 hr earlier than what has been observed during infection of HeLa cells^41^ and correlates with the differential *O. tsutsugamushi* growth and *ank5* expression kinetics in these cell types (Extended Data Figure 3B–3D).

**Figure 3.**
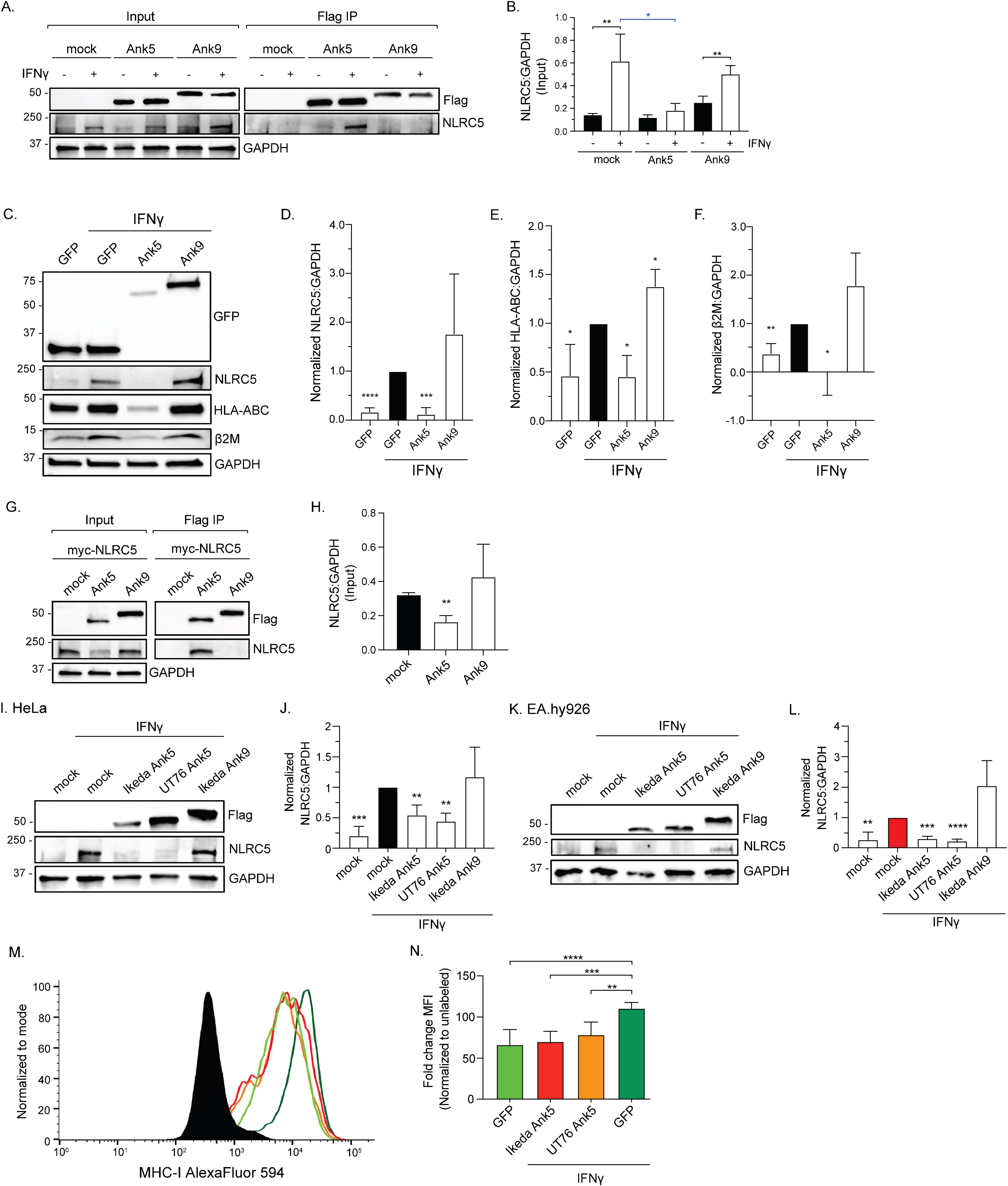
Ank5 decreases levels of NLRC5, cellular levels of MHC class I components, and surface levels of MHC class I. (A and B) HeLa cells were mock transfected or transfected to express Flag-Ank5 or Flag-Ank9 and treated with vehicle [-] or IFNγ [+]. (A) “Input” lysates and immunoprecipitated Flag-tag protein complexes “Flag IP” were analyzed by immunoblotting. (B) Densitometric quantification of NLRC5:GAPDH signal from input blots. Data are means ± SD (n = 4). An unpaired *t*-test determined statistical significance among pairs. Indicators of statistical significance between pairs of untreated (black bars) and IFNγ-treated (white bars) samples are colored black. Indicators of statistical significance between transfected and IFNγ-treated samples compared to mock transfected and IFNγ-treated samples are colored blue. (C–F) HeLa cells were transfected to express indicated proteins, treated with IFNγ or vehicle control and cell sorted on GFP-positivity. (C) Whole cell lysates were analyzed by immunoblotting. (D–F) The densitometric signal value of NLRC5 (D), HLA-ABC (E), or β2M (F) was divided by the corresponding GAPDH densitometric signal value and normalized to the densitometric signal level of GFP-expressing cells that had been treated with IFNγ (black bars). Data are means ± SD (n = 3). An unpaired *t*-test determined statistical significance among pairs compared to GFP-expressing cells treated with IFNγ (black bars). (G and H) HeLa cells were mock transfected or transfected to co-express Flag-Ank5 or Flag-Ank9 with myc-NLRC5. (G) Input lysates and immunoprecipitated Flag-tag protein (Flag IP) complexes were analyzed by immunoblotting. (H) Densitometric quantification of the NLRC5:GAPDH signal ratio from input blots. Data are means ± SD (n = 3). An unpaired *t*-test determined statistical significance among pairs compared to mock transfected samples (black bars). (I–L) HeLa (I and J) or EA.hy926 (K and L) cells were mock transfected or transfected to express Flag-tagged Ikeda Ank5, UT76 Ank5, or Ikeda Ank9 and treated with IFNγ. (I and K) Whole cell lysates were analyzed by immunoblotting. (J and L) Densitometric quantification of the NLRC5:GAPDH signal ratios from input blots. Data are means ± SD (n = 3). An unpaired *t-*test was used to determine statistical significance among pairs compared to mock transfected cells (black bar, HeLa; red bar, EA.hy926). (M and N) Surface levels of MHC class I from sorted cells were analyzed by flow cytometry. (M) Representative histograms of MHC class I levels normalized to mode. Unlabeled (black) samples served as the control. Colors coordinate to those used in (N). (N) The mean fluorescence intensity of each sample was divided by that of its unlabeled control per replicate. Data are means ± SD (n = 7). One-way ANOVA with Tukey’s *post hoc* test was used to assess significant differences between every mean. *, p < 0.05; **, p < 0.01; ***, p < 0.001; ****, p < 0.0001.

To assess if Ank5-mediated reduction of NLRC5 inhibits expression of MHC class I components, HeLa cells were transfected to express GFP-Ank5, GFP-Ank9, or GFP and treated with IFNγ. Fluorescence-activated cell sorting (FACS) was used to enrich for GFP-positive cells. GFP-Ank5 but not GFP or GFP-Ank9 decreased levels of NLRC5, HLA-ABC, and β2M (Figure 3C–3F). The slight elevation of HLA-ABC levels in GFP-Ank9-expressing cells could be due to the inhibitory effect by Ank9 on the secretory pathway.^64^ When myc-NLRC5 was co-expressed with GFP-tagged Ank5, Ank9, Ank4 (inhibits endoplasmic reticulum-associated degradation^66^), Ank6 (blocks NF-κB accumulation in the nucleus^55,65^), or Ank13 (nucleomodulin^67^), its levels were significantly lower exclusively in cells expressing GFP-Ank5 (Extended Data Figure 4E and 4F). GFP-Ank5 also specifically co-precipitated and reduced levels of myc-NLRC5 (Figure 3G and 3H). When UT76 Ank5 was ectopically expressed in IFNγ stimulated HeLa or EA.hy926 cells, it functioned similarly to Ikeda Ank5 by reducing NLRC5 and cell surface MHC class I levels (Figure 3I to 3N). Thus, ectopically expressed Ikeda and UT76 Ank5 proteins can bind and reduce cellular levels of NLRC5, which lowers amounts of MHC class I proteins to recapitulate phenomena associated with *O. tsutsugamushi* infection.

### Ank5 directs ubiquitination of NLRC5 at K1194 and its 26S proteasomal degradation in an F-box-dependent manner

NLRC5 is degraded by the 26S proteasome in *O. tsutsugamushi* infected cells. To test if Ank5-mediated reduction of NLRC5 is also proteasome-dependent, HeLa cells expressing myc-NLRC5 with or without Flag-Ank5 were treated with MG132. In mock transfected cells, proteasome inhibition resulted in higher levels of myc-NLRC5 (Figure 4A). Because cells were transfected to express Flag-Ank5 prior to the addition of MG132 or vehicle, NLRC5 levels were considerably lower regardless of treatment. Nevertheless, Flag-Ank5 coprecipitated three-fold more NLRC5 in the presence of MG132 (Figure 4B), indicating an interaction exists between Ank5 and NLRC5 that promotes 26S proteasomal degradation of the transactivator.

**Figure 4.**
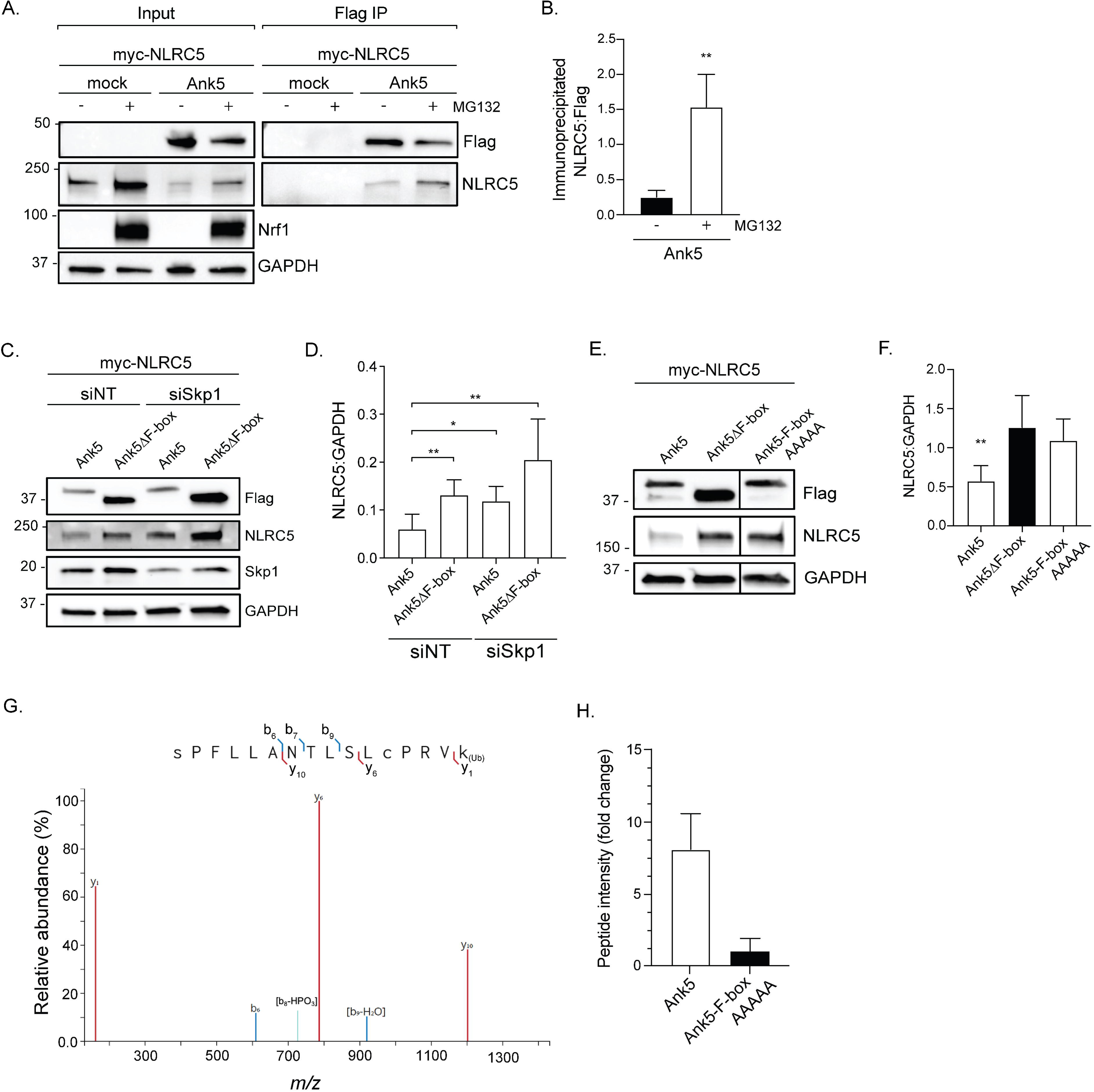
Ank5 directs ubiquitination and proteasomal degradation of NLRC5 in an F-box-dependent manner. (A and B) HeLa cells were transfected to express myc-NLRC5 alone or together with Flag-Ank5 followed by the addition of vehicle [-] or MG132 [+]. (A) Both input lysates and immunoprecipitated Flag-tag protein (Flag IP) complexes were analyzed by immunoblotting. (B) The densitometric signal value of immunoprecipitated NLRC5 was divided by that of the corresponding Flag-Ank5 signal. Data are means ± SD (n = 3). An unpaired *t*-test was used to determine statistical significance among pairs. (C and D) HeLa cells were transfected with nontargeting (siNT) or Skp1-targeting (siSkp1) siRNA. The cells were then transfected to co-express myc-NLRC5 and either Flag-Ank5 or Flag-Ank5ΔF-box. (C) Whole cell lysates were analyzed by immunoblotting. (D) Densitometric quantification of the NLRC5:GAPDH signal ratio. Data are means ± SD (n = 5). An unpaired *t*-test was used to determine statistical significance among pairs. (E and F) HeLa cells were transfected to co-express myc-NLRC5 with indicated Flag-Ank5 proteins. (E) Whole cell lysates were analyzed by immunoblotting. Vertical lines between bands indicate where the blot was cropped or imaged separately. (F) Densitometric quantification of NLRC5:GAPDH signal. Data are means ± SD (n = 6). An unpaired *t-*test was used to determine statistical significance among pairs compared to Flag-Ank5ΔF-box (black bar). (G and H) HeLa cells transfected to co-express Flag-NLRC5 and either GFP-Ank5 or GFP-Ank5-F-boxAAAAA were treated with MG132 and cell sorted on GFP-positivity. Flag-NLRC5 was immunoprecipitated from collected lysates, eluted, and the eluted complexes were analyzed by mass spectrometry. (G) Fragmentation spectra of the post-translationally modified _1179_SPFLLANTLSLCPRVK_1194_ peptide. Spectra-matched b- (blue) and y-ion (red) fragments are displayed and summarized by the trypsinized peptide sequence inset above. (H) Peptide counts presented as mean fold change ± SEM (n = 2). *, p < 0.05; **, p < 0.01.

The Ank5 F-box domain binds Skp1 to assemble the SCF E3 ubiquitin ligase complex.^55^ To confirm if the F-box and Skp1 contribute to the effector’s ability to reduce NLRC5 cellular levels, HeLa cells that had been treated with Skp1 targeting small interfering RNA (siRNA) or non-targeting siRNA were transfected to co-express myc-NLRC5 and either Flag-Ank5 or Flag-Ank5ΔF-box. Flag-Ank5 failed to degrade myc-NLRC5 in Skp1 knockdown cells (Figure 4C and 4D). Also, myc-NLRC5 levels remained elevated in cells expressing Flag-Ank5ΔF-box whether Skp1 had been knocked down or not. As another means of validating the essentiality of the F-box for Ank5 to direct NLRC5 degradation, HeLa cells were transfected to express myc-NLRC5 together with Flag-Ank5ΔF-box or Flag-tagged Ank5 having F-box residues L301, P302, E304, I309, and D317 that are required for binding Skp1 and nucleating the SCF complex replaced with alanine (Flag-Ank5-F-boxAAAAA).^55^ Flag-Ank5ΔF-box and Flag-Ank5-F-boxAAAAA were similarly incapacitated for the ability to degrade NLRC5 (Figure 4E and 4F).

To delineate the NLRC5 lysine(s) that Ank5 directs to become ubiquitinated, HeLa cells were transfected to co-express Flag-NLRC5 and GFP-tagged Ank5 or Ank5-F-boxAAAAA. The cells were treated with MG132 and GFP positive populations enriched for via FACS. Lysates were verified for expression of proteins of interest and MG132 treatment (Extended Data Figure 5). LC-MS/MS analysis of precipitated Flag-NLRC5 revealed that the same NLRC5 peptide (_1179_SPFLLANTLSLCPRVK_1194_) that is ubiquitinated at K1194 in *O. tsutsugamushi* infected cells (Figure 1J) is ubiquitinated at K1194 in cells expressing GFP-Ank5 (Figure 4G), though the overall amount was lower than in infected cells, explaining the fewer product ions observed. That said, this peptide was recovered at seven-fold greater abundance from cells expressing GFP-Ank5 compared to cells expressing GFP-Ank5-F-boxAAAAA (Figure 4H). Altogether, these data confirm that ectopically expressed Ank5 phenocopies *O. tsutsugamushi* infection by directing SCF ubiquitination of NLRC5 at K1194 and its proteasomal degradation in an F-box-dependent manner.

### Ank5 AR four is critical for binding NLRC5

Guided by the principle that AR domains mediate protein-protein interactions,^82-84^ we sought to identify the specific Ank5 AR(s) that bind NLRC5. Because individual ARs are modular and as long as a minimum of two repeats is maintained for stability, deletion of a single repeat within an AR domain does not significantly alter tertiary structure of the domain or the entire protein.^85-87^ Accordingly, plasmids were generated that encoded Flag-Ank5-F-boxAAAAA proteins lacking each of the four ARs to allow for assessment of the ΔAR domain-NLRC5 interaction without the complication of NLRC5 being degraded. AlphaFold predicted that all of the Ank5 ARs are tandemly arranged helix-turn-helix motifs that form a concave interaction interface consistent with known AR domain structure^82,88^ (Extended Data Figure 6). The F-box of each was predicted to adopt a fold prototypical of other F-box containing proteins.^55,89-91^ Importantly, when the ΔAR mutant models were overlaid on full length Ank5-F-boxAAAAA, no structural alterations were predicted for any ΔAR protein’s AR domain (Extended Data Figure 6). Flag-Ank5-F-boxAAAAA and the ΔAR1 and ΔAR2 versions thereof co-immunoprecipitated myc-NLRC5 with comparable efficiencies (Figure 5A and 5B). The ability of Ank5-F-boxAAAAA to co-immunoprecipitate NLRC5 was impaired when AR3 was deleted and abolished when AR4 was removed. UT76 Ank5 AR4 and Ikeda Ank5 AR4 are identical (Extended Data Table 3). Whereas Flag-tagged UT76 Ank5ΔF-box precipitated NLRC5, Flag-tagged UT76 Ank5ΔAR4 could not (Extended Data Figure 7), indicating that, like Ikeda Ank5, UT76 Ank5 requires its AR4 to bind NLRC5 and its F-box to direct NLRC5 degradation.

**Figure 5.**
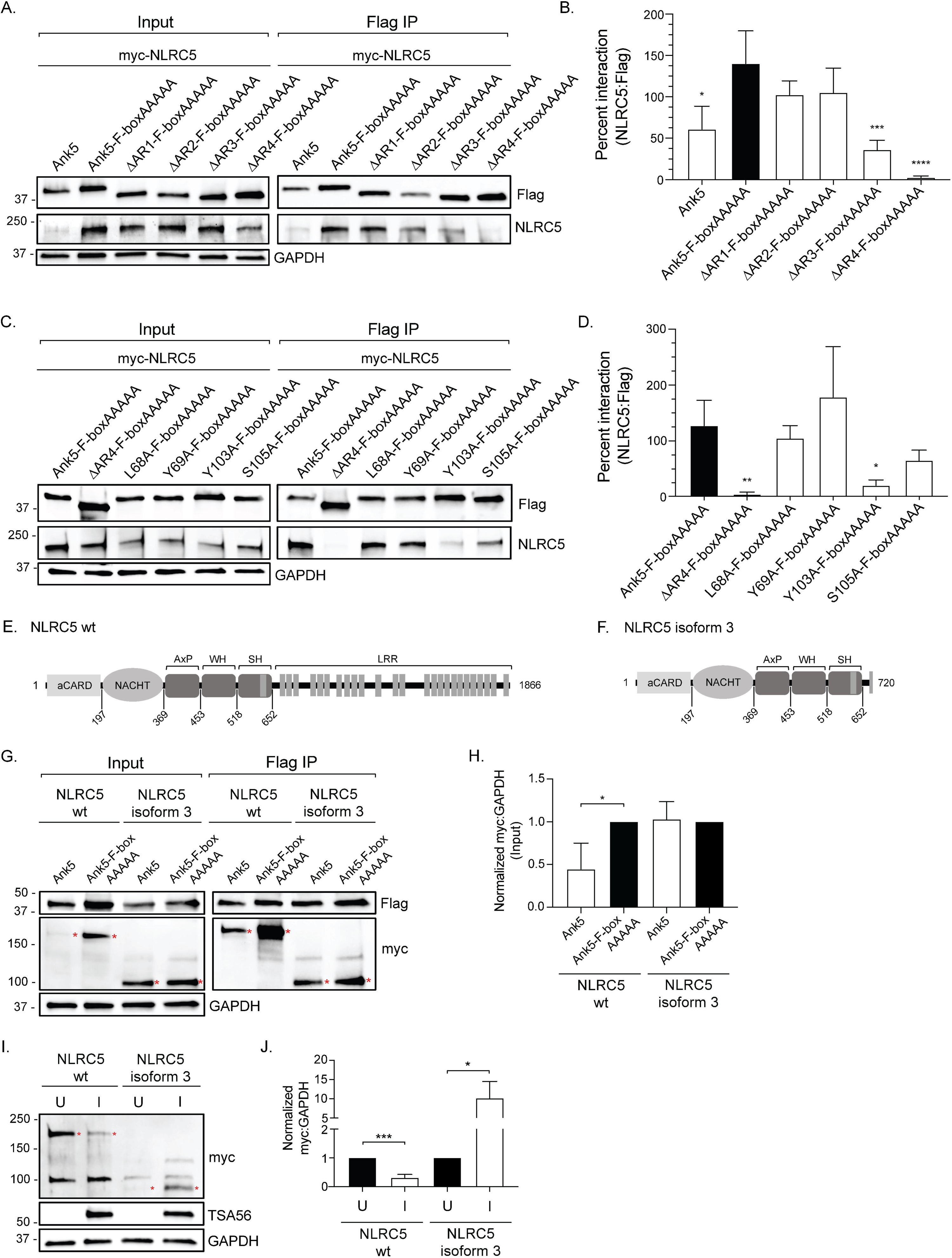
Ank5 AR4 is critical for binding NLRC5 and the NLRC5 LRR is essential for Ank5-mediated degradation. (A–D) HeLa cells were transfected to co-express myc-NLRC5 and Flag-tagged Ank5, Ank5-F-boxAAAAA, ΔAR-F-boxAAAAA (ΔAR AAAAA) proteins, or F-boxAAAAA proteins bearing the indicated additional alanine substitution. (A and C) Both input lysates and immunoprecipitated Flag-tag protein (Flag IP) complexes were analyzed by immunoblotting. (B and D) NLRC5 densitometric values were divided by the corresponding Flag densitometric signal. The quotients were multiplied by 100 to determine percent interaction. Data are means ± SD (n = 3). An unpaired *t-*test was used to determine statistical significance among pairs compared to Flag-Ank5-F-boxAAAAA (black bar). (E and F) Schematics of wt NLRC5 (E) and isoform 3 (F). The atypical CARD (aCARD), NACHT, small helical (AxP), winged helix (WH), and superhelical (SH) domains of each are denoted. Wt NLRC5 contains a complete leucine rich repeat (LRR) region consisting of 26 tandemly arrayed LRRs (E), whereas isoform 3 lacks this region (F). Amino acid positions at the beginning of each domain are indicated. (G–J) HeLa cells were transfected to co-express myc-tagged wt NLRC5 or isoform 3 with either Flag-tagged Ank5 or Ank5-F-boxAAAAA (G and H) or transfected to express myc-tagged wt NLRC5 or isoform 3 and then mock infected [U] or infected with *O. tsutsugamushi* [I] (I and J). (G and I) Input lysates, immunoprecipitated Flag-tag protein (Flag IP) complexes, and whole cell lysates were analyzed by immunoblotting. Red asterisks denote the true bands of interest. (H and J) Densitometric quantification of the NLRC5:GAPDH signal ratios from input blots. Data are means ± SD (n = 3). An unpaired *t*-test determined statistical significance among pairs compared to Flag-Ank5-F-boxAAAAA (H) or uninfected (J) samples. *, p < 0.05; **, p < 0.01; ***, p < 0.001, ****, p < 0.0001.

The β-hairpin and first α-helix of each 33-residue AR create the recognition surface. Within this region, conserved residues preserve tertiary structure while non-conserved or semi-conserved residues at positions 2, 3, and 5 in the β-turn mediate protein-protein interactions and dictate target specificity.^82,89,92^ In agreement with pulldown studies (Figure 5A and 5B), NovaDock predicted that Ank5 AR3 and AR4 bind NLRC5. Of the eight candidate interacting residues, two in AR3 (L68 and Y69) and two in AR4 (Y103 and S105) correspond to AR3 positions 2 and 3 and AR4 positions 3 and 5 (Extended Data Figure 8). We investigated the contribution of these four amino acids to NLRC5 binding. The remaining four could not be evaluated because they occupy positions within the inner α-helix that would alter AR tertiary structure if mutated.^93-96^ Flag-Ank5-F-boxAAAAA proteins with the residues of interest substituted with alanine were assessed for the ability to bind myc-NLRC5 compared to the positive control, Flag-Ank5-F-boxAAAAA, and negative control, Flag-Ank5ΔAR4-F-boxAAAAA. Changing L68 or Y69 to alanine had no effect on the interaction while swapping S105 for alanine yielded a detectable but statistically insignificant reduction in NLRC5 recovery (Figure 5C and 5D). The ability of the Y103A mutant to co-immunoprecipitate NLRC5 was nearly abolished. Thus, Ank5 requires AR4 to bind NLRC5, and this interaction critically involves Y103.

### NLRC5 residues 1 to 720 are essential for Ank5 binding and its LRR is required for Ank5 to promote its degradation

NLRC5 is a multi-domain protein.^97-99^ Its N-terminal aCARD (<u>a</u>typical <u>c</u>aspase <u>a</u>ctivation and <u>r</u>ecruitment <u>d</u>omain) is followed by NACHT (present in <u>NA</u>IP, <u>C</u>IITA, <u>H</u>ET-E, and <u>T</u>P1), small helical (AxP), winged helix (WH), and superhelical (SH) domains.^99^ Its C-terminal half consists of 26 tandemly arranged LRRs. There are six NLRC5 isoforms with varying LRR region lengths that result from differential splicing.^40^ Relative to full-length NLRC5 (Figure 5E), isoform 3 consists only of residues 1 to 720 and lacks the entire LRR domain (Figure 5F). Because LRR regions can mediate protein-protein interactions,^100^ we assessed the relevance of the NLRC5 LRR region in the Ank5 interaction by comparing the abilities of Flag-Ank5 and Flag-Ank5-F-boxAAAAA to co-immunoprecipitate myc-tagged versions of wild-type (wt) NLRC5 and isoform 3. A myc tag antibody was used to detect myc-NLRC5 proteins because the NLRC5 antibody used thus far recognizes a C-terminal epitope that isoform 3 lacks. The myc antibody detected a nonspecific host cell protein in all samples (Figure 5G). Flag-Ank5 and Flag-Ank5-F-boxAAAAA precipitated both isoforms revealing that the LRR region is dispensable for Ank5 to bind NLRC5 proteins. Interestingly, Flag-Ank5 was unable to degrade myc-NLRC5 isoform 3 (Figure 5G and 5H), insinuating that the LRR region is necessary for Ank5-mediated NLRC5 degradation. When cells expressing myc-tagged wt NLRC5 or isoform 3 were infected with *O. tsutsugamushi*, wt NLRC5 levels were severely reduced whereas isoform 3 levels were significantly elevated (Figure 5I and 5J). Therefore, *O. tsutsugamushi* Ank5 binds to NLRC5 within residues 1 to 720 but requires the C-terminal LRR domain to orchestrate its degradation.

### Ank5 inhibits MHC class I expression in an AR4-dependent but F-box-independent manner

The ability of Ank5 to degrade NLRC5 is AR4- and F-box-dependent. Yet, it was unclear if Ank5 requires both domains to impair MHC class I expression. HeLa cells expressing GFP, GFP-Ank5, GFP-Ank5-F-boxAAAAA, or GFP-Ank5ΔAR4 were treated with IFNγ to upregulate NLRC5 expression, sorted on GFP positivity to enrich for each transfected population, and measured for NLRC5 and MHC class I mRNA and protein expression. Compared to unstimulated controls, IFNγ treated cells exhibited significantly elevated NLRC5 and MHC class I transcript and protein levels as well as MHC class I cell surface amounts (Figure 6). *NLRC5* mRNA levels were unchanged in cells expressing GFP-Ank5, GFP-Ank5-F-boxAAAAA, and GFP-Ank5ΔAR4 compared to cells expressing GFP (Figure 6A). Expression of GFP-tagged Ank5 or Ank5-F-boxAAAAA, but not Ank5ΔAR4, resulted in lower transcript levels of *HLA-A*, the first gene in the HLA locus,^101,102^ and *β2M* despite GFP-Ank5 being the only one of the three proteins that promoted NLRC5 degradation (Figure 6B–6E). Similarly, HLA-ABC and β2M total protein along with MHC class I cell surface levels were significantly reduced by GFP-Ank5 and GFP-Ank5-F-boxAAAAA but not GFP-Ank5ΔAR4 (Figure 6F–6I). Overall, these data indicate that Ank5 binding NLRC5 by virtue of AR4 is sufficient to impair MHC I gene expression, which translates to losses in overall and cell surface levels of MHC class I.

**Figure 6.**
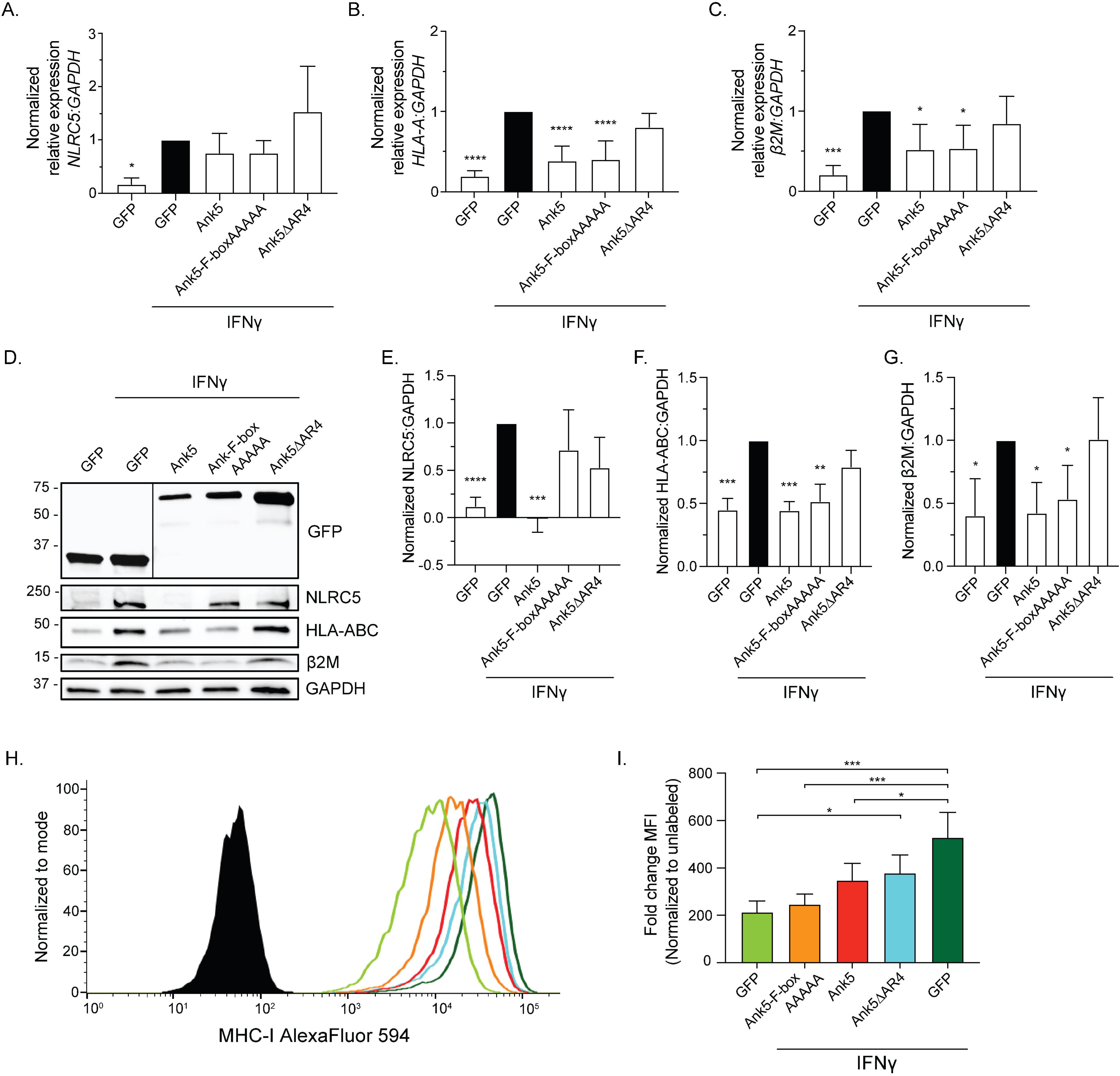
Ank5 inhibits MHC class I expression in an AR4-dependent but F-box-independent manner. (A–I) HeLa cells transfected to express indicated proteins were treated with IFNγ or vehicle control and sorted on GFP-positivity. (A–C) Total RNA was collected and RT-qPCR was performed using gene-specific primers. Relative expression of *NLRC5*-to-*GAPDH* (A), *HLA-A*-to-*GAPDH* (B), and *β2M*-to-*GAPDH* (C) was determined using the 2^-ΔΔCT^ method in which conditional values were normalized to those values of GFP-expressing cells that had been treated with IFNγ (black bars). One-way ANOVA with Dunnett’s *post hoc* test was used to assess significant differences where every mean value was compared the mean value of GFP-expressing cells that had been treated with IFNγ (black bar). Data are means ± SD (n = 5). (D–G) Whole cell lysates were collected from sorted cells. (D) Whole cell lysates were analyzed by immunoblotting. Vertical lines between bands indicate where the blot was cropped or imaged separately. (E–G) The densitometric signal value of NLRC5 (E), HLA-ABC (F), or β2M (G) was divided by the corresponding GAPDH densitometric signal value and normalized to the densitometric signal level of GFP-expressing cells that had been treated with IFNγ (black bars). Data are means ± SD (n = 3). An unpaired *t-*test was used to determine statistical significance among pairs compared to GFP-expressing cells that had been treated with IFNγ (black bars). (H and I) Surface levels of MHC class I from sorted cells was analyzed by flow cytometry. (H) Representative histograms of MHC class I levels normalized to mode. Unlabeled (black) samples served as the control. Colors coordinate to those used in (I). (I) The mean fluorescence intensity of each sample was divided by that of its unlabeled control per replicate. Data are means ± SD (n = 4). One-way ANOVA with Tukey’s *post hoc* test was used to assess significant differences between every mean. *, p < 0.05; **, p < 0.01; ***, p < 0.001; ****, p < 0.0001.

### Ectopically expressed Ank5-F-boxAAAAA competitively antagonizes *O. tsutsugamushi* mediated NLRC5 degradation

To confirm whether bacterial derived Ank5 promotes NLRC5 degradation in infected cells, we needed an approach that would circumvent *O. tsutsugamushi* genetic intractability. Since ectopically expressed Ank5 engages NLRC5 via AR4 and promotes its degradation using its F-box, we rationalized that overexpressed Ank5-F-boxAAAAA binding to NLRC5 would competitively antagonize the actions of *Orientia*-derived Ank5. HeLa cells were transfected to express GFP-tagged Ank5, Ank5-F-boxAAAAA, Ank5ΔAR4, Ank5ΔAR4-F-boxAAAAA or GFP. Each sorted population was infected with *O. tsutsugamushi* or mock infected followed by the addition of IFNγ. At 24 hr p.i., endogenous NLRC5 levels were significantly lower in infected versus uninfected cells expressing GFP and comparably reduced in uninfected and infected cells expressing GFP-Ank5 (Figure 7). NLRC5 amounts were also significantly lower in cells expressing GFP-Ank5ΔAR4 and GFP-Ank5ΔAR4-F-boxAAAAA during *O. tsutsugamushi* infection, indicating the inability of either protein to outcompete bacterial derived Ank5. However, NLRC5 levels were unchanged between uninfected and infected cells expressing GFP-Ank5-F-boxAAAAA, validating the ability of the ectopically expressed protein to protect NLRC5 from degradation mediated by oriential Ank5. These results also verify that Ank5 expressed by *O. tsutsugamushi* during infection binds NLRC5 via its AR4 to direct its degradation in an F-box-dependent manner.

**Figure 7.**
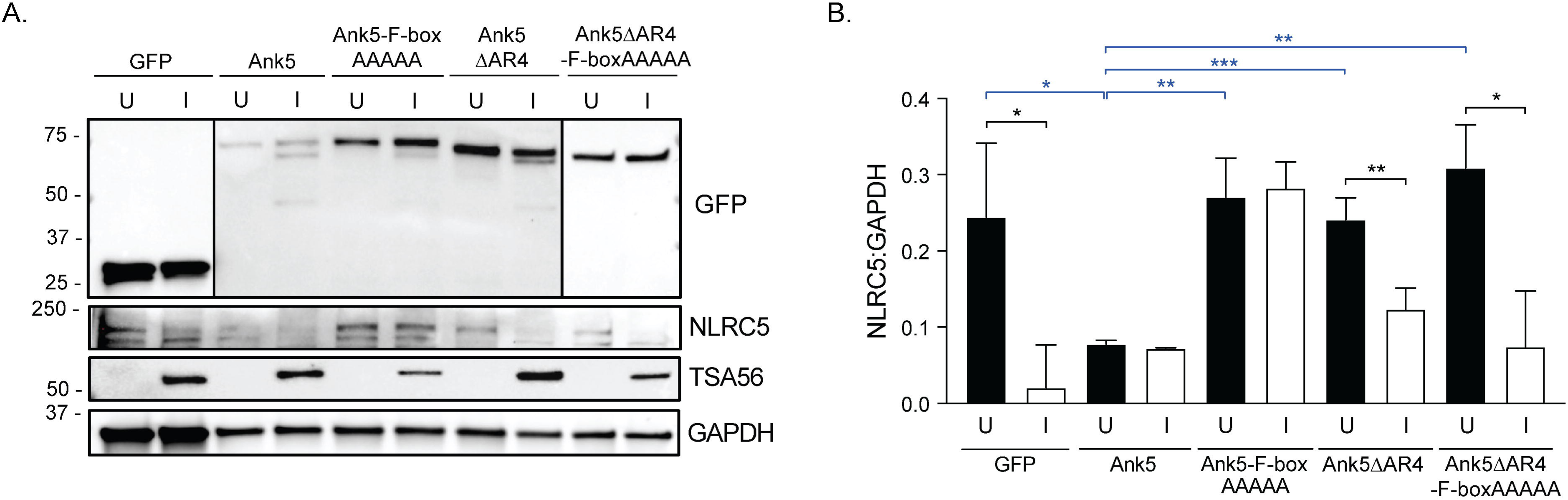
Ank5-F-boxAAAAA competitively antagonizes *O. tsutsugamushi* mediated NLRC5 degradation. (A and B) HeLa cells transfected to express the indicated proteins were sorted on GFP positivity, mock infected [U] or infected with *O. tsutsugamushi* [I] and treated with IFNγ. (A) Whole cell lysates were analyzed by immunoblotting. Vertical lines between bands indicate where the blot was cropped or imaged separately. (B) Densitometric quantification of the NLRC5:GAPDH signal ratios. Data are means ± SD (n = 3). An unpaired *t-*test was used to determine statistical significance among pairs. Indicators of statistical significance between pairs of uninfected (black bars) and infected (white bars) samples are colored black. Indicators of statistical significance between transfected and uninfected cells compared to uninfected cells expressing GFP-Ank5 are colored blue. *, p < 0.05; **, p < 0.01; ***, p < 0.001.

## DISCUSSION

Like *O. tsutsugamushi*, many other intracellular microbes deploy AR-F-box proteins that counter eukaryotic immunity, benefit microbial fitness, and drive pathogenicity.^53,57-62^ Yet nearly all these proteins’ modes of action are unknown. Here we comprehensively elucidated the mechanism by which *O. tsutsugamushi* Ank5 binds NLRC5 and directs its ubiquitination and proteasomal degradation to impair MHC class I expression. In doing so, we reveal NLRC5 as a novel target for microbial immunomodulation. Prior to this report, known strategies by which viruses and intracellular bacteria subvert the MHC class I pathway included altering subcellular transport of, inhibiting peptide loading on, and increasing degradation of MHC class I molecules.^2-4,6,7,103^ *Mycobacterium tuberculosis* protein PPE38 modestly reduces MHC class I transcription when expressed in *Mycobacterium smegmatis*, but the underlying mechanism has not been examined.^5^ *O. tsutsugamushi* directs ubiquitination of NLRC5 on K1194 and promotes its degradation in a MG132-sensitive manner. Ectopically expressed Ank5 phenocopies both effects. Ank5-mediated NLRC5 degradation translates to decreased *HLA-A* and *β2M* transcription, which reduces overall and cell surface MHC class I levels. The ability of Ank5 to direct NLRC5 ubiquitination and proteasomal degradation requires its F-box and host Skp1, findings that are consistent with the observation that Ank5 interactions with Skp1 and two other SCF complex proteins, Cul1 and RING box 1, are F-box-dependent.^55^ Overexpression of dominant negative Ank5-F-boxAAAAA that can bind but not degrade NLRC5 protects it from *Orientia*-induced degradation by outcompeting bacterial-derived Ank5. Overall, Ank5 is a primary effector for *O. tsutsugamushi* modulation of NLRC5-dependent MHC class I expression.

Ank5 binds the N-terminal half of NLRC5. This interaction requires Ank5 AR4 and critically involves Y103 therein. The absolute sequence conservation of AR4 among Ikeda, UT76, and Wuj/2014 Ank5 proteins suggests that its functional essentiality has been maintained by selective pressure. This contention is further supported by the conserved abilities of Ikeda and UT76 to lower NLRC5 and MHC class I cellular levels during infection and for ectopically expressed Ikeda Ank5 and UT76 Ank5 to recapitulate these phenomena in an AR4- and F-box-dependent manner. NLRC5 and its differential splice variants are expressed in hematopoietic cells and tissues. Whereas isoform 3 ends at residue 720 and lacks the LRR domain, the other five splice variants extend beyond K1194 and have LRR regions.^40^ Ank5 binds wt NLRC5 and isoform 3 but cannot promote degradation of isoform 3, an observation that reinforces the importance of K1194 and links the LRR region to NLRC5 susceptibility to Ank5-mediated ubiquitination and proteasomal degradation. Because the LRR region is critical for MHC class I transactivation,^104^ this finding also supports that *O. tsutsugamushi* strains that express Ank5 would be able to rid their host cells of all NLRC5 isoforms competent for transactivating MHC class I. *O. tsutsugamushi* promotes a more complex NLRC5 interactome than in uninfected cells. Aside from those involved in protein degradation, the roles of proteins in this interactome in mediating NLRC5 degradation or other modulatory functions remain to be determined.

NLRC5 deficiency has profound impacts on adaptive immunity. In *nlrc5*^-/-^ mice, MHC class I surface expression is severely reduced in CD4^+^ T cells, CD8^+^ T cells, and B cells.^105-107^ APCs from these mice are impaired for inducing MHC class I-dependent antigen-specific stimulation of co-cultured CD8^+^ T cells for proliferation, IFNγ production, and cytolytic activity.^105,106,108^ Consequently, *nlrc5*^-/-^ mice are more susceptible to intracellular pathogens as they exhibit higher *Listeria monocytogenes* loads and are less effective at clearing influenza A and rotavirus infections.^106-109^ NLRC5 elevated expression in cancer cells is associated with MHC class I gene expression, susceptibility to CD8^+^ T-cell-mediated killing, and tumor growth inhibition.^110-113^ Conversely, its suppression correlates with reduced MHC class I expression and lower cancer patient survival rates. Immune escape of cancer cells due to NLRC5 suppression can result from multiple mechanisms.^110^ However, the mode of action by which the proto-oncogene BMI1 impairs the NLRC5-MHC class I axis is of notable mention because it is reminiscent of Ank5. BMI1, an E3 ubiquitin ligase component of polycomb repressive complex 1, binds NLRC5 to induce its ubiquitination and degradation, which reduces MHC class I surface expression and CD8^+^ T cell activation.^114^

Of more than 1900 bacterial genomes, the Ikeda chromosome encodes the fifth-largest number of Anks, accounting for 2.4% of its content. By comparison, *ank* genomic content for *Homo sapiens*, *Mus musculus*, and the average of all obligate intracellular bacteria except *Chlamydia* species, which do not have any *ank* genes, is 3.1%, 1.7%, and 0.8%, respectively.^51^ Retention of such a high *ank* content over its reductive evolution suggests the importance of Anks to *O. tsutsugamushi* pathobiology. However, very few have been studied and their functions are poorly characterized.^64-67,69^ The significance of Ank5 to *O. tsutsugamushi* pathogenesis is apparent. By upregulating Ank5 during cytoplasmic replication in endothelial cells, the bacterium can orchestrate NLRC5 degradation so that it can grow to high numbers in host cells rendered incapable of eliciting adaptive immunity via MHC class I antigen presentation. The *Leptotromidium* mites that vector *O. tsutsugamushi* are considered its primary reservoirs.^10^ However, laboratory-conducted studies demonstrated that naïve chiggers could acquire the bacterium from infected mice and in rare instances their offspring could infect new hosts.^115,116^ Suppressing the NLRC5-MHC class I axis could augment *Orientia* reaching a sufficient load in rodent hosts that maximizes its chance for reacquisition via mite feeding before immune responses capable of clearing the infection are elicited. Given that mites lack adaptive immunity, the Ank5 mechanism uncovered here is exclusively relevant to *Orientia* infection in mammals and argues that mammals might play a role in shaping the immunomodulatory/virulence capacity of *O. tsutsugamushi* strains. Ank5-mediated immunosuppression would also potentially contribute to disease outcomes associated with infection of endothelial cells in accidental human hosts. High *O. tsutsugamushi* burdens are observed during late disease, and severe manifestations in patients and experimentally infected mice at least partially stem from endothelial cell colonization.^31,117-123^

Differences in genomic carriage and expression of Ank5 could influence scrub typhus pathogenesis. Indeed, while NLRC5 and MHC class I levels are pronouncedly reduced in cells infected with Ikeda or UT76, NLRC5 expression is markedly increased and MHC class I levels unaltered during Karp infection. Therefore, Ank5 affords strains that carry and express it the advantage of downregulating NLRC5-dependent anti-oriential immune responses. Notably, the studies that concluded that MHC class I and CD8^+^ T cells are vital for preventing fatal *O. tsutsugamushi* infections and CD8^+^ T cells contribute to dysregulated Th1 immunopathologies in murine models were conducted using Karp.^32,33,117,120^ Yet, despite being unable to degrade NLRC5 and block HLA-ABC and β2M expression, Karp still impairs MHC class I cell surface presentation. Although the responsible mechanism is undefined, Karp might execute this strategy using Ank9, which inhibits the secretory pathway.^48,52,64^

NLRC5 has been implicated in signal transduction, Type I interferon responses, selective autophagy, and possibly inflammasome activation.^124-126^ *O. tsutsugamushi* strains that carry/express Ank5 or other Anks could contribute to strain-specific variations in scrub typhus immunopathologies independent of MHC class I. In support of this premise, a dual RNA-seq comparison of endothelial cells infected with Karp or *O. tsutsugamushi* strain UT176 showed that UT176 induced a stronger proinflammatory response and was cleared in mice while Karp stimulated IL-33 alarmin-based immunopathologies and was pathogenic in mice. Although neither strain carries *ank*5, they differentially express putative virulence factors during infection, including several *anks.*^24^ Given that *O. tsutsugamushi* Anks varies strain-to-strain,^42,48,49,52^ and their functions are mostly unknown, future work must define additional modes of action for these effectors and determine their roles in scrub typhus pathogenesis.

In conclusion, our study establishes Ank5 as an *O. tsutsugamushi* virulence factor that promotes NLRC5 degradation using two eukaryotic-like motifs that simultaneously bind NLRC5 and nucleate the SCF ubiquitin ligase complex. As an adaptor protein in this complex scaffold, Ank5 directs selective ubiquitination of NLRC5 on K1194 to target it to the proteasome. In the absence of NLRC5, MHC class I gene expression, total protein, and surface levels are significantly reduced. The remarkable similarity between Ank5 and BMI1 mechanisms of action is a striking example of convergent evolution in that both an intracellular pathogen and oncogene co-opt ubiquitination/proteasomal degradation to subvert the NLRC5-MHC class I axis. As exemplified for BMI1,^114^ this strategy likely enables Ikeda and other *O. tsutsugamushi* strains that express Ank5 to evade CD8+ T cell-mediated immunity and additional NLRC5-dependent immune responses. By unveiling NLRC5 as a target for microbial immunomodulation, this study advances understanding of how *O. tsutsugamushi* and potentially other intracellular pathogens maximize survival in their intracellular niches.

## METHODS

### Cultivation of cell lines and *O. tsutsugamushi* infections

Uninfected HeLa cells (CCL-2; American Type Culture Collection [ATCC], Manassas, VA) and HeLa cells infected with *O. tsutsugamushi* str. Ikeda (NC_010793.1) were maintained as previously described.^65^ EA.hy926 endothelial cells (CRL-2922; ATCC) were cultured in Dulbecco’s modified Eagle’s medium (DMEM) with l-glutamine, 4.5 g d-glucose and 100 mg sodium pyruvate (Gibco, Gaithersburg, MD) supplemented with 10% (vol/vol) heat-inactivated fetal bovine serum (FBS) (Gemini Bioproducts, West Sacramento, CA), 1X minimal essential medium containing non-essential amino acids (Gibco) and 15 mM HEPES (Gibco) in a humidified incubator at 37 °C with 5% atmospheric CO_2_. The Ikeda, UT76, and Karp strains were used in this study. For routine propagation, bacteria were grown in HeLa cells. For experimental use, infected HeLa cells (≥ 90%) were mechanically disrupted using glass beads followed by differential centrifugation to recover host cell-free bacteria as described^65^ and then subsequently incubated with naïve HeLa or EA.hy926 cells. Cells undergoing mock infections were incubated with an equivalent volume of bead-lysed HeLa or EA.hy926 cells. Synchronous infections were achieved by replacing inoculum with fresh media at 2 to 4 hr post infection (p.i.) Experiments were verified for achieving a multiplicity of infection (MOI) of 10 by assessing coverslips of infected cells by immunofluorescence as described below.

### Plasmid constructs

Empty pEGFP-C1 was included as a control.^50^ pFlag-Ank4, -Ank5, -Ank6, -Ank9, -Ank13, -Ank5ΔF-box, pGFP-Ank5^50^, and pFlag-Ank5-F-boxAAAAA^55^ were generated previously.^50^ pFlag-Ank5-F-boxAAAAA was also generated previously.^55^ Primers used to introduce the following mutations (Supplementary Table 1) were designed using the In-Fusion Cloning Primer Design Tool v1.0 (www.takarabio.com). pFlag-Ank5ΔAR1-F-boxAAAAA (lacks residues 3-33), -Ank5ΔAR2-F-boxAAAAA (lacks residues 34-66), -Ank5ΔAR3-F-boxAAAAA (lacks residues 67-100), -Ank5ΔAR4-F-boxAAAAA (lacks residues 101-133), pFlag-Ank5_L68A_-F-boxAAAAA, pFlag-Ank5_Y69A_-F-boxAAAAA, pFlag-Ank5_Y103A_-F-boxAAAAA, and pFlag-Ank5_Y105A_-F-boxAAAAA were generated using TaKaRa Bio USA (San Francisco, CA) In-Fusion Mutagenesis protocol and pFlag-Ank5-F-boxAAAAA as template. pFlag-Ank5ΔAR4 and pGFP-Ank5ΔAR4 were generated using In-Fusion Mutagenesis and pFlag-Ank5 or pGFP-Ank5 as template, respectively. pGFP-Ank5-F-boxAAAAA and GFP-Ank5ΔAR4-F-boxAAAAA were generated by digesting pFlag-Ank5-F-boxAAAAA and pFlag-Ank5ΔAR4-F-boxAAAAA with EcoRI-HF and BamHI and subcloning the released fragments into the multicloning site in pEGFP-C1. Mammalian codon optimized UT76 *ank5* was synthesized and cloned into p3xFlag-CMV-7.1 downstream and in-frame with a 3xFlag tag coding sequence by Genewiz (Burlington, MA) to generate pFlag-UT76-Ank5. pFlag-UT76-Ank5 served as template for the generation of pFlag-UT76-Ank5ΔF-box and pFlag-UT76-Ank5ΔAR4 using TaKaRa Bio USA (San Francisco, CA) In-Fusion Mutagenesis protocol and previously described Ank5ΔF-box^55^ and herein designed Ank5ΔAR4 primers. pCMV-Tag2b-Flag-NLRC5 (Addgene plasmid #37521; http://n2t.net/addgene:37521; RRID:Addgene_37521), pcDNA3.1-3xmyc-B-NLRC5 (Addgene plasmid #37509; http://n2t.net/addgene:37509; RRID:Addgene_37509), and pcDNA3.1-3xmyc-B-NLRC5 iso3 (Addgene plasmid #37510; http://n2t.net/addgene:37510; RRID:Addgene_37510) were gifts from Thomas Kufer. pcDNA3.1-HA-Ubiquitin (Ub) was a gift from Edward Yeh (Addgene plasmid #18712; http://n2t.net/addgene:18712; RRID:Addgene_18712). Full length *tsa56* (OTT_0945) was amplified from *O. tsutsugamushi* str. Ikeda genomic DNA with *tsa56*-Full-F and *tsa56*-Full-R primers. The subsequent amplicon was cloned into TOPO vector pCR4.0 to generate pCR4.0::*Ot_tsa56*. All plasmid constructs were confirmed by sequence analysis (Genewiz).

### Transfection

HeLa or EA.hy926 cells grown to 90% confluency were transfected with plasmid DNA using Lipofectamine 2000 (Invitrogen, Carlsbad, CA) and incubated at 37 °C in a humidified incubator at 5% atmospheric CO_2_ for 18 to 24 h. The amount of DNA used for transfections was modified from that recommended by the Lipofectamine 2000 protocol per plasmid to accommodate varying levels of transfection efficiency, as determined by western blot densitometric signal. After incubation, spent media was removed. The cells were washed once with PBS before being processed for further applications.

### siRNA knockdown

HeLa cells grown to 90% confluency were transfected with ON-TARGETplus human Skp1 SMARTpool siRNA or non-targeting siRNA (GE Healthcare Dharmacon, Inc., Lafayette, CO) at a final concentration appropriate to the vessel size. After 48 hr, the media was replaced and knocked down cells were transfected to express either Flag-Ank5 or Flag-Ank5ΔF-box. After 24 hr, lysates were collected and western-blotted as described above.

### Pharmacologic and IFNγ treatments

To prevent eukaryotic protein synthesis or proteasome degradation during infection, *O. tsutsugamushi-* or mock-infected HeLa cells were treated with or without cycloheximide (50 μg ml^-1^ in ethanol; Sigma-Aldrich) or MG132 (5 μM in DMSO; Sigma-Aldrich), respectively, at 2 hr p.i. and whole cell lysates were collected at 18 hr p.i. To inhibit the proteasome in transfected cells, cells were first transfected for 18 to 24 hr and then treated with 5 μM MG132 or DMSO for 18 to 24 hr prior to collection. To increase endogenous NLRC5 levels, at 24 hr after transfection or 2 hr p.i., spent media was replaced with media containing 40 ng ml^-1^ human IFNγ (PeproTech, Rock Hill, NJ) or vehicle control (0.1% bovine serum albumin [BSA] in H_2_O) until collection.

### Immunofluorescence

Cells were fixed and permeabilized with -20°C methanol. MOI coverslips and flow cytometry coverslips were blocked in 5% (vol/vol) bovine serum albumin (BSA) in phosphate buffered saline (PBS; 1.05 mM KH_2_PO_4_, 155 mM NaCl, 2.96 mM Na_2_HPO_4_, pH 7.4) then incubated with rabbit anti-TSA56 (^64^; 1:1,000) followed by incubation with Alexa Fluor 488-conjugated goat anti-rabbit ([A11034], 1:10,000) or Alexa Fluor 594-conjugated goat anti-rabbit ([A11012], 1:10,000) in 5% BSA. Blocking and antibody incubations were performed for 1 hr at room temperature with three PBS washes between each step. MOI coverslips were incubated with 300 nM 4’ 6-diamidino-2-phenylindole (DAPI; [D1306] Invitrogen) in PBS for 5 min, washed three times, and mounted using ProLong Gold Anti-fade reagent (Invitrogen). MOI coverslips were imaged, and percentage of infection was calculated with an Olympus BX51 spinning disc confocal microscope (Olympus, Shinjuku City, Tokyo, Japan). Flow cytometry coverslip images were obtained using a Zeiss LSM 700 laser scanning confocal microscope (Zeiss).

### Immunoprecipitation

For Fab-trap immunoprecipitations, HeLa cells expressing HA-Ub and Flag-NLRC5 for 18 hr were synchronously infected with *O. tsutsugamushi* str. Ikeda as described above. At 24 hr p.i., cells were incubated with MG132 for 24 hr followed by lysis in high saline Tris buffer (50 mM Tris HCl, 400 mM NaCl, 1 mM EDTA, pH 7.4) with 1.0% Triton x-100 (TBHS-T) spiked with Halt Protease and Phosphatase Inhibitor Cocktail (100X; Thermo). ChromoTek DYKDDDDK Fab-trap Agarose resin (Proteintech Group, Inc., Rosemont, IL) was washed with TBHS-T buffer two times, centrifuged at 2,500 x g for two min, and added to normalized cell lysates in a final volume of 400 μl. Samples were incubated with beads rotating at 4 °C for 1 hr followed by centrifugation at 2,500 x g for 5 min and washing with TBHS-T four times. Proteins were eluted with Laemmli buffer containing 4% βME and a fraction of the eluate was resolved by SDS-PAGE next to input lysates and immunoblotted as described below.

For anti-Flag affinity agarose immunoprecipitations, transfected HeLa cells were harvested and lysed in TBHS-T spiked with Halt Protease and Phosphatase Inhibitor Cocktail. Protein A/G agarose beads (ThermoFisher Scientific) were washed with TBHS-T buffer three times, centrifuged at 8,600 x g for 30 s, and added to normalized cell lysates in a final volume of 400 μl. The samples were rotated with beads at 4 °C for 4 hr followed by centrifugation at 8,600 x g for 30 s. Recovered supernatants were added to Anti-Flag M2 Affinity Gel (MilliporeSigma) that had been washed three times with TBHS-T buffer. Samples rotated with beads at 4 °C overnight followed by centrifugation at 8,600 x g for 30 s and washed with TBHS-T six to ten times. Washed beads were resuspended in Laemmli buffer containing 4% βME and incubated at 100 °C for 5 min to elute bound proteins. Inputs (10-30 μg) and eluates were resolved by SDS-PAGE and screened by western blot.

### Western blotting

Western blotting was performed as previously described.^67^ Briefly, normalized amounts of eluates were resolved by SDS-PAGE in 4 to 15% TGX polyacrylamide gels (Bio-Rad) at 110 V for 15 min followed by 200 V for 25 min. Proteins were transferred onto nitrocellulose membrane in Towbin buffer at 100 V for 30 min. Blots were blocked in either 5% (vol/vol) non-fat dry milk or 5% (vol/vol) BSA in tris-buffered saline plus 0.05% Tween 20 (TBS-T) after which they were screened with rat anti-NLRC5 (MilliporeSigma [MABF260], 1:1,000), rabbit anti-TSA56^64^ (1:1,000), rabbit or mouse anti-Flag (Sigma-Aldrich [F7425 or F1804], 1:1,000), rabbit anti-GFP (ThermoFisher Scientific, Rockford, IL [A-6455], 1:1,000), mouse anti-GAPDH (Santa Cruz, Dallas, TX [sc-365062], 1:750), rabbit or mouse anti-myc (Abcam, Cambridge, United Kingdom [ab9106 and ab32], 1:1,000), rabbit anti-HA (Abcam [ab9110], 1:4,000), mouse anti-HLA-ABC heavy chain (Abcam [ab70328], 1:1,000), rabbit anti-Β2M (Life Technologies, Carlsbad, CA [4H5L5], 1:1,000), rabbit anti-Skp1 (Cell Signaling Technology, Danvers, MA [2156S], 1:750), or rabbit anti-Nrf1 (Cell Signaling Technology [D5B10], 1:1,000). Bound primary antibodies were detected using horseradish peroxidase (HRP)-conjugated horse anti-mouse, anti-rabbit, and anti-rat IgG (Cell Signaling Technology, [7076S, 7074S, 7077S], 1:10,000). All blots were incubated with either SuperSignal West Pico PLUS, SuperSignal West Dura, or SuperSignal West Femto chemiluminescent substrate (ThermoFisher Scientific) prior to imaging in a ChemiDoc Touch Imaging System (Bio-Rad). Bio-Rad Image Lab 6.0 software was used to obtain densitometric values.

### Mass spectrometry-based immunoprecipitation proteomics

HeLa cells were transfected to express Flag-NLRC5 for 18 hr and then synchronously infected with *O. tsutsugamushi* str. Ikeda as described above. At 24 hr p.i., cells were incubated with 5 μM MG132 for 24 hr. In parallel, HeLa cells were transfected to express Flag-NLRC5 with either GFP-Ank5 or GFP-Ank5-F-boxAAAAA for 18 hr after which point all samples were incubated with MG132 for 24 hr. Populations of GFP-positive cells were isolated using FACS as described below and lysed with TBHS-T spiked with Halt Protease and Phosphatase Inhibitor Cocktail. Flag-NLRC5 was immunoprecipitated using Fab-trap agarose resin. The beads were washed, resuspended in elution buffer (65 mM Tris HCl, 2% SDS, with 50 mM dithiothreitol (DTT)), and incubated at 100 °C for 10 min to elute bound proteins. Inputs (10-30 μg) were resolved by SDS-PAGE and screened by western blot. Eluates were stored at -80 °C until processing for mass spectrometry. Eluates were load balanced (20 µg) and surfactant depleted with Pierce Detergent Removal Spin Columns (ThermoFisher Scientific). Samples were reduced with TCEP (10 mM), carboxymethylated with iodoacetamide (15 mM), and quenched with DTT (10 mM). In solution digests were performed overnight at pH 8 and 37 °C with mass spectrometry grade trypsin (Promega, Madison, WI) at a 1:33 enzyme-to-protein ratio and quenched with formic acid (0.15%). Peptide digests were analyzed as described previously.^127^ Samples were loaded onto a Symmetry C18 trap column on a NanoAcquity (Waters, Milford, MA) ultra-performance liquid chromatography system and gradient eluted from 6% to 44% acetonitrile in water (0.1% formic acid) over 90 min at 400 nL min^-1^ using a 150 mm x 75 µm HSS T3 column maintained at 55 °C. Eluting peptides were electrosprayed into a Synapt G2-Si tandem mass spectrometer (Waters). Data were acquired in data-independent acquisition mode with ion-mobility separation and collision energy optimization (UDMSe). Ions were resolved between 400 and 1800 m/z at a nominal resolution of 25,000. Spectra were peak picked, integrated and precursor-to-product aligned using PLGS 3.0.3 with data aligned across biological replicates by retention time (± 0.8 min), mobility drift time (± 2 bins) and precursor ion mass (MH^+^, ± 8 ppm) with EndogeSeq. Data were filtered to ion features that replicated in over 50% of samples and searched against the tryptic peptides of the human NLRC5 protein (accession Q86WI3) and matched to within ± 4 ppm precursor ion mass and ± 6 ppm product ion mass while accounting for fixed carboxymethylation and variable ubiquitination and phosphorylation with results controlled to a 5% false discovery rate. Ion peak area counts for ubiquitinated peptides were median centered, imputed for the limit of quantification, log-2 transformed and assessed as fold-change from respective control conditions with a student’s *t*-test.

### Yeast two-hybrid

ULTimate yeast two-hybrid analysis was performed by Hybrigenics Services (Paris, France). Mammalian codon optimized *O. tsutsugamushi* str. Ikeda *ank5ΔF-box* was PCR amplified and cloned as an *N*-terminal LexA fusion into pB27 (*N*-*lexA*-*ank5ΔF-box*-*C*). Sequence fidelity was confirmed, and the construct was introduced into yeast (bait) and screened through mating with yeast bearing a randomly primed human placental library (prey). Positively selected clones were isolated, and captured prey fragments were PCR amplified, sequenced, and identified using NCBI Protein Basic Local Alignment Search Tool (BLASTP) (https://blast.ncbi.nlm.nih.gov/Blast.cgi). The probability of specificity for each interaction was calculated using a predicted biological score (PBS), described as very high (A), high (B), or good (C) confidence between the two proteins.^128^

### Analysis of *ank5* homologs

The National Center for Biotechnology Information (NCBI) Nucleotide Basic Local Alignment Search Tool (BLASTN) was used to identify homologs of the representative Ikeda *ank5* copy (OTT_RS01000) present in the genomes of other *O. tsutsugamushi* strains in GenBank. BLASTN analyzed nucleotide sequence identity and BLASTP analyzed amino acid identity of Ikeda Ank5 homologs identified in UT76 (UT76HP_01926) and Wuj/2014 (CP044031).

### qPCR

Total genomic DNA (gDNA) was isolated using the DNeasy Mini Kit (Qiagen) and eluted in 20 to 30 μl of AE buffer. Concentration and purity were determined using a spectrophotometer (NanoVue Plus; GE, Boston, MA). Serial dilutions were prepared using pCR4.0::*Ot_tsa56* to generate a standard curve. qPCR of standards and normalized gDNA was performed with PerfeCTa SYBR Green FastMix (QuantaBio, Beverly, MA) and primer pairs against *tsa56* (Supplementary Table 1). Thermal cycling conditions used were 95 °C for 30 s, followed by 40 cycles of 95 °C for 5 s and 54 °C for 15 s. *O. tsutsugamushi* genome equivalents (GE) were quantified based on the standard curve using the CFX Maestro for Mac 1.0 software package (Bio-Rad).

### RT-qPCR

Total RNA was isolated using the RNeasy Mini Kit (Qiagen) and eluted in 25 μl of RNase-free water. Concentration and purity were determined. One μg of RNA was treated with amplification grade DNase I (Invitrogen) following the manufacturer’s protocol. cDNA was generated using the iScript Reverse Transcription Supermix protocol (Bio-Rad). Parallel reactions were performed in the absence of reverse transcriptase. To verify successful genomic DNA depletion, both samples were subjected to PCR with human *GAPDH*-specific primers^41^ and MyTaq polymerase (Bioline, Taunton, MA). Resulting PCR amplicons were visualized by agarose gel electrophoresis. qPCR using cDNA as template was performed with SsoFast EvaGreen supermix (Bio-Rad) and primer pairs against *O. tsutsugamushi* 16S rDNA (*ott16S*)^66^ or *ank5.*^50^ Primers against *ank5* were designed to produce amplicons from any one of the three identical *ank5* copies. Thermal cycling conditions used were 95 °C for 30 s, followed by 40 cycles of 95 °C for 5 s and 55 °C for 5 s. Relative expression was determined using the 2^−ΔΔCT^ method^129^ as part of the CFX Maestro for Mac 1.0 software package (Bio-Rad).

### FACS

HeLa cells were washed with PBS, trypsinized, and recovered by centrifugation at 500 x g for 5 min. The resulting cell pellets were resuspended in 1 ml of pre-sort buffer (EDTA (Versene) Solution 0.526 mM (Irvine Scientific, Santa Ana, CA) with 2% (vol/vol) heat-inactivated FBS) and flushed through a 40 µm mesh cell strainer (Greiner Bio-One, Monroe, NC). GFP-expressing cells were sorted from non-transfected cells on a Becton Dickinson (BD) FACSMelody using BDFACSChorusTM 1.3.3 (BD Biosciences) in Purity Sort mode. The GFP population was gated on positivity compared to non-transfected HeLa cells. Sorted cells were collected in 10% (vol/vol) FBS in PBS and pelleted at 500 x g for 5 min before progressing to additional assays.

### Use of ectopically expressed Ank5 proteins to competitively antagonize *O. tsutsugamushi* derived Ank5

HeLa cells were transfected to express GFP-Ank5, GFP-Ank5-F-boxAAAAA, GFP-Ank5ΔAR4, GFP-Ank5ΔAR4-F-box-AAAAA, or GFP. At 18 hr, the cells were subjected to FACS to isolate GFP-positive cells. Recovered cells were centrifugated at 500 x g for 5 min, resuspended in 1 ml RPMI containing 10% FBS, and halved into 6-well plates. Plates were spun at 1,000 x g for 3 min to optimize cell adherence and incubated in a humidified incubator at 35 °C with 5% atmospheric CO_2_ for 18 h. Next, the cells were infected with *O. tsutsugamushi* str. Ikeda as described above. At 2 hr p.i., media was replaced with fresh RPMI containing 1% (vol/vol) FBS and 40 ng ml^-1^ human IFNγ for 24 h. At this point, cells were collected, washed, lysed, and western-blotted as described above.

### Flow cytometry

Transfected or *O. tsutsugamushi* infected HeLa cells were washed with PBS and lifted from the flask by incubating in EDTA (Versene) Solution 0.526 mM for 10 min. Transfected cells were pelleted and processed for FACS as described above. Recovered cells were incubated with Fc block (Miltenyi Biotec, Bergusch Gladbach, Germany, 1:100), vortexed, then incubated for 5 min on ice. A portion of the Fc blocked-cells were separated to accommodate the unlabeled control while remaining samples were incubated with mouse anti-human HLA-ABC W6/32 (Invitrogen [MA5-11723], 1:40) on ice for 30 min. Between each step, cells underwent centrifugation at 1.5 rpm for 5 min and washed with 1 ml pre-sort buffer or PBS. Cells were incubated with Alexa Fluor 594-conjugated goat anti-mouse IgG (Invitrogen [A11032], 1:100) for 20 min on ice followed by fixation with 500 μl of 4% (vol/vol) paraformaldehyde (Electron Microscopy Services, Hatfield, PA) on ice for 20 min. Cells were washed once more with 1 ml PBS and resuspended in 300 μl PBS before flow cytometry was performed on a BD FACSMelody using BDFACSChorusTM 1.3.3. Ten thousand cells were analyzed per sample using FlowJo (version 10.8.1) software (BD Biosciences).

### Ank5-NLRC5 protein-protein interaction prediction

NovaFold (DNASTAR, Madison, WI), which is based on the I-TASSER algorithm,^130-132^ was used to model the NLRC5 NACHT domain using iterative assembly simulations. NLRC5 is 1866 amino acids in length, which exceeds the allowable size limit for NovaDock interaction modeling. Because the LRR region was found to be dispensable for the Ank5-NLRC5 interaction herein, we utilized the NACHT domain in modeling studies. Our rationale was further guided by the shared protein architecture between NLRC5 and a related NLR protein, CIITA. CIITA and NLRC5 both bind the host cell ankyrin protein, RFXANK, as part of the transactivating enhanceosome^35^ and CIITA utilizes its NACHT domain to do so,^133,134^ suggesting this region of NLRC5 could be bound by Ank5. NovaDock (DNASTAR) predicted atomic protein docking interactions between Ank5 and the NLRC5 NACHT domain using the SwarmDock algorithm.^135^ Protean 3D (DNASTAR) was used to visually depict the NovaDock predictions.

### Bioinformatic analysis

STRING^75^ was used to analyze interaction networks. AlphaFold^136,137^ was used to predict tertiary structures for Ank5-F-boxAAAAA, Ank5ΔAR1-F-boxAAAAA, Ank5ΔAR2-F-boxAAAAA, Ank5ΔAR3-F-boxAAAAA, and Ank5ΔAR4-F-boxAAAAA. Alignment and color coding of the resulting program database files were done using the PyMOL Molecular Graphic System, version 1.2r3pre, Schrödinger, LLC.

### Statistical analysis

Statistical analyses were performed using the Prism 8.0 software package (GraphPad, San Diego, CA). Unpaired student *t*-test was used to assess differences between pairs. One-way ANOVA with Tukey’s *post hoc* test was used to test for a significant difference among group means. One-way ANOVA with Dunnett’s *post hoc* test was used to test for a significant difference among group means compared to a control group mean. Statistical significance was set at *P*-values of < 0.05.

## AUTHOR CONTRIBUTIONS

Conceptualization, H.E.A. and J.A.C.; investigation, H.E.A, J.R.H., A.K.O., K.G.R.; writing – original draft, H.E.A. and J.A.C.; writing – review and editing, H.E.A., A.K.O., J.A.C.; supervision, J.A.C.; acquisition of grant support, J.A.C., A.K.O., and H.E.A.

## ACKNOWLEDGEMENTS

We would like to thank Dr. Rebecca Martin for assisting with flow cytometry data analysis and Dr. Paige E. Allen, Dr. Travis J. Chiarelli, Dr. Erol Fikrig, Dr. Leigh A. Knodler, Dr. Curtis B. Read, Dr. Jeanne Salje, Dr. Savannah E. Sanchez, and Dr. Mary M. Weber for critical review of this manuscript. This work was supported by National Institutes of Health-National Institute of Allergy and Infectious Diseases (www.niaid.gov) grants 1R01 AI123346, 2R56 AI123346, and 1R01 AI167857 to J.A.C., and American Heart Association grant 20PRE35210610 to H.E.A. Laser scanning confocal was performed at the VCU Microscopy Facility, which is funded, in part, by the NIH-NCI Cancer Center Support Grant P30 CA016059.

## Summary Model

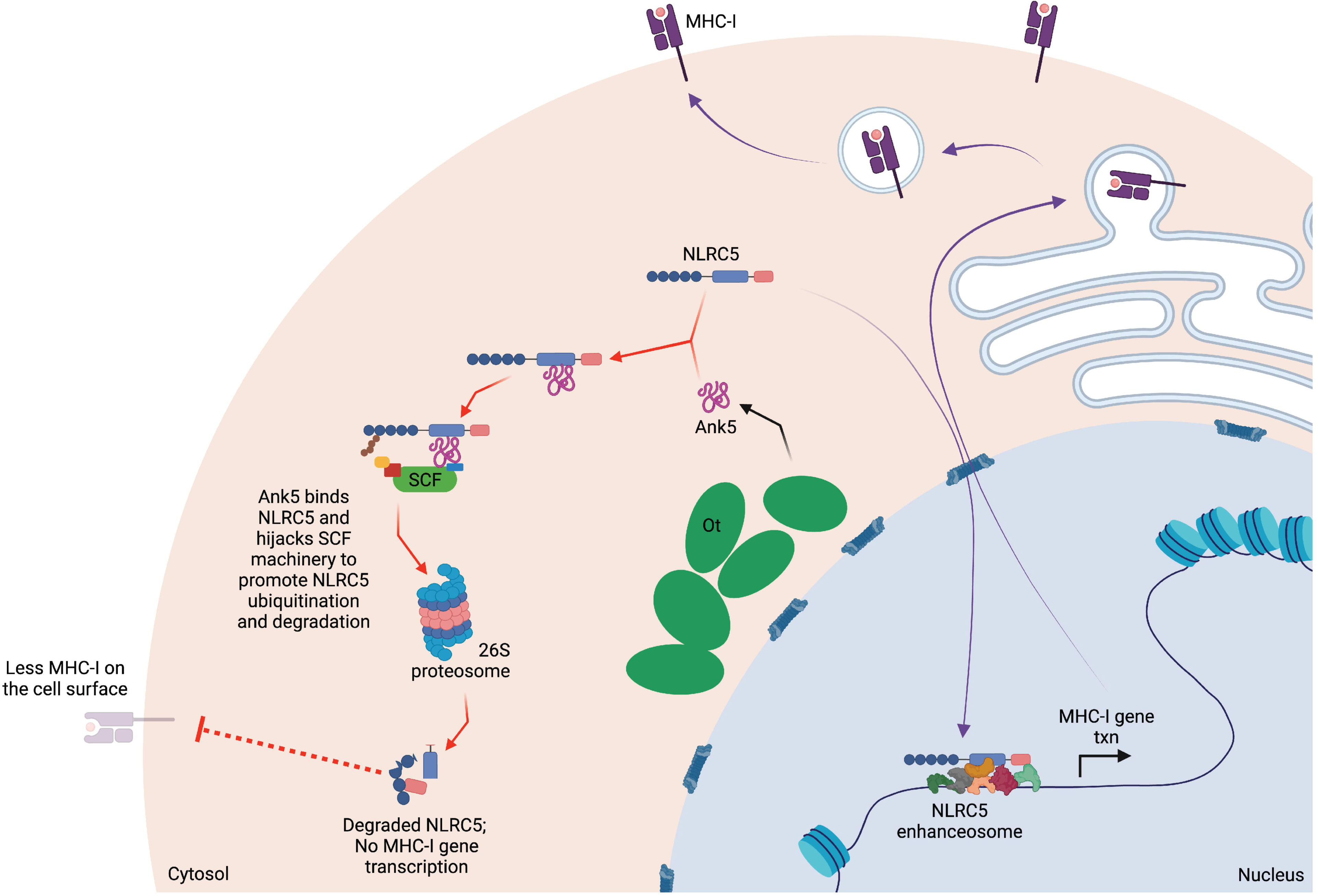

## FIGURE LEGENDS

**Extended Data Figure 1.**
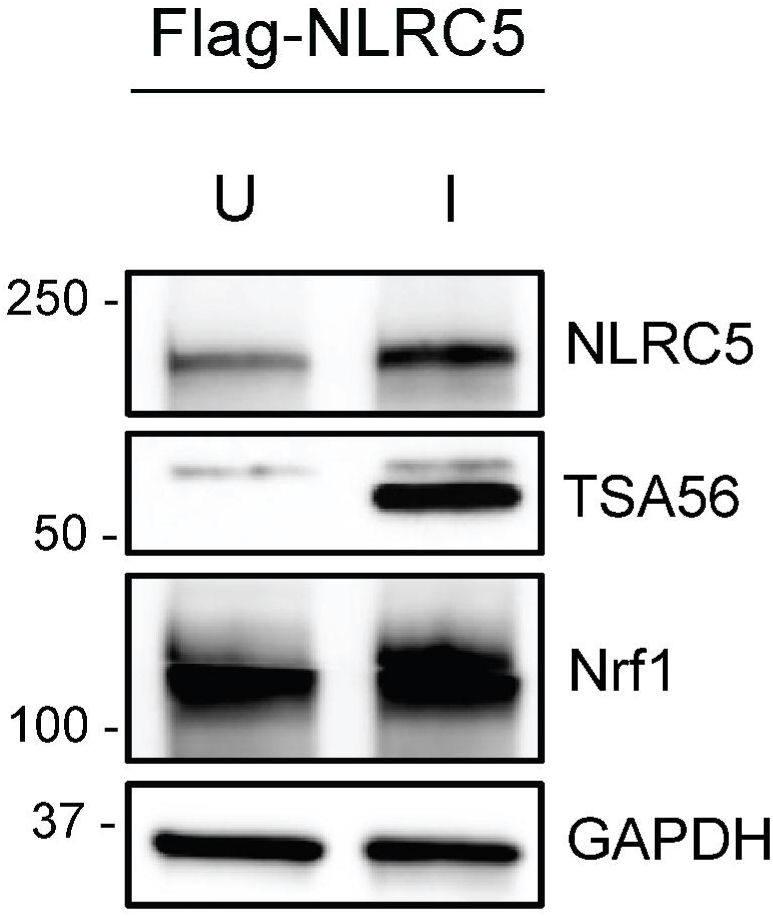
Input lysates from uninfected and *O. tsutsugamushi*-infected cells for LC-MS/MS IP-Proteomics. Western blots of input lysates for LC-MS/MS IP-proteomics to verify successful MG132 treatment and expression of all proteins of interest by cells that were transfected to express Flag-NLRC5 and then either mock infected [U] or infected with *O. tsutsugamushi* [I].

**Extended Data Figure 2.**
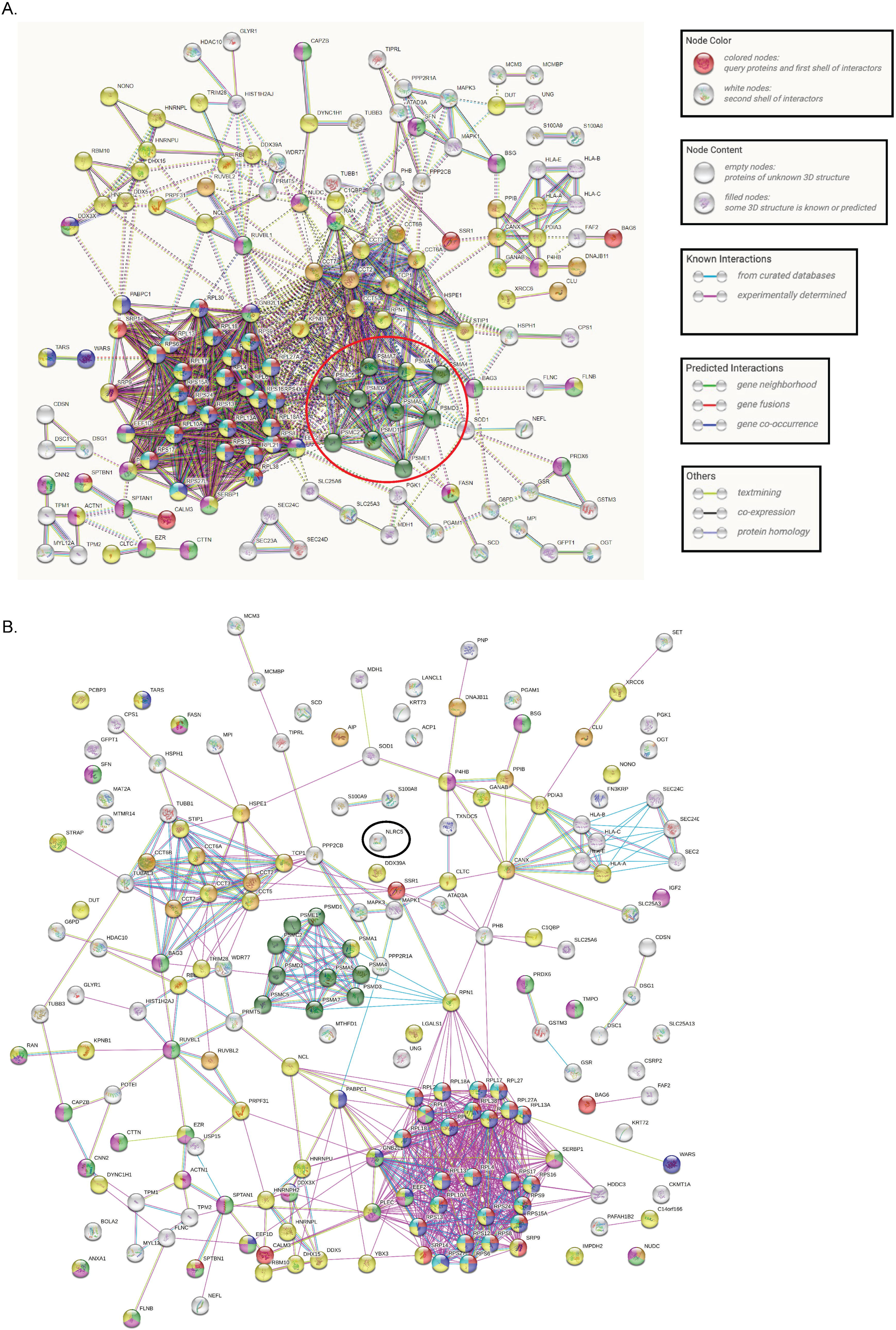
NLRC5 interactome recovered from *O. tsutsugamushi* infected cells. STRING analysis of the interactome co-immunoprecipitated by Flag-NLRC5 from cells infected with *O. tsutsugamushi* relative to uninfected cells. (A) The red circle was manually added to denote proteasomal subunit proteins. (B) NLRC5 (denoted by black circle) was manually added to the interactome described in (A) and then re-analyzed to only display known physical interaction networks.

**Extended Data Figure 3.**
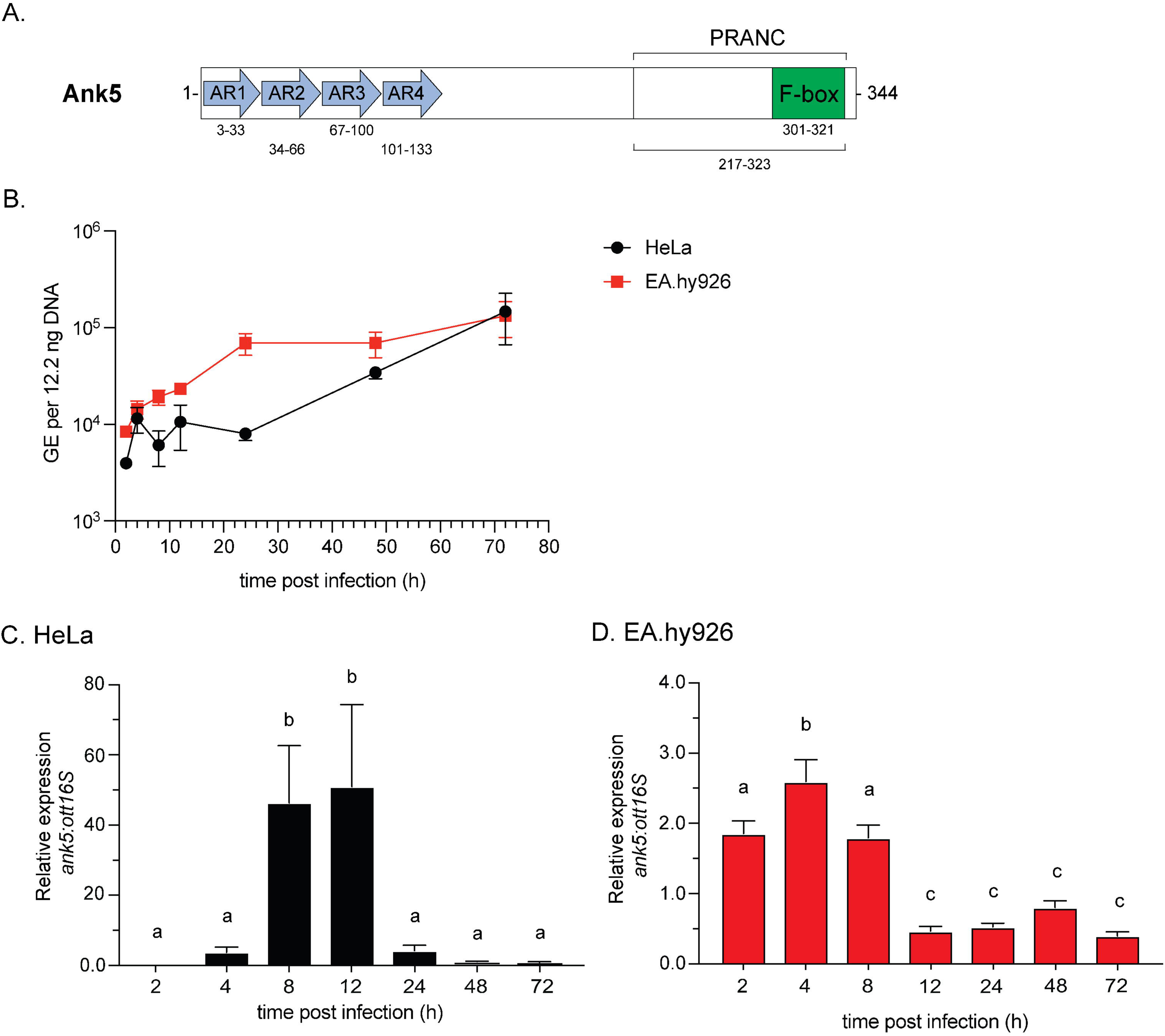
*O. tsutsugamushi* transcriptionally upregulates *ank5* prior to expansive growth. (A) Schematic of Ank5 depicting its four tandemly arranged ARs (blue arrows) and F-box (green). The F-box occurs as part of a larger encompassing PRANC domain (white). Amino acids that comprise each domain are indicated. (B) *O. tsutsugamushi* replication was measured during infection of HeLa and EA.hy926 cells via quantification of *O. tsutsugamushi* genomic equivalents (GE) over the course of infection. Data are means ± SEM (n = 3). (C and D) Relative expression of *O. tsutsugamushi ank5*-to-16S rRNA gene (*ott16S*) during infection of HeLa (C) and EA.hy926 (D) cells was determined using the 2^-ΔΔCT^ method. Data are means ± SD (n = 3). One-way ANOVA with Tukey’s *post hoc* test was used to test for significant difference of relative *ank5* levels across time points. Mean values labeled a, b, and c are significantly different from each other.

**Extended Data Figure 4.**
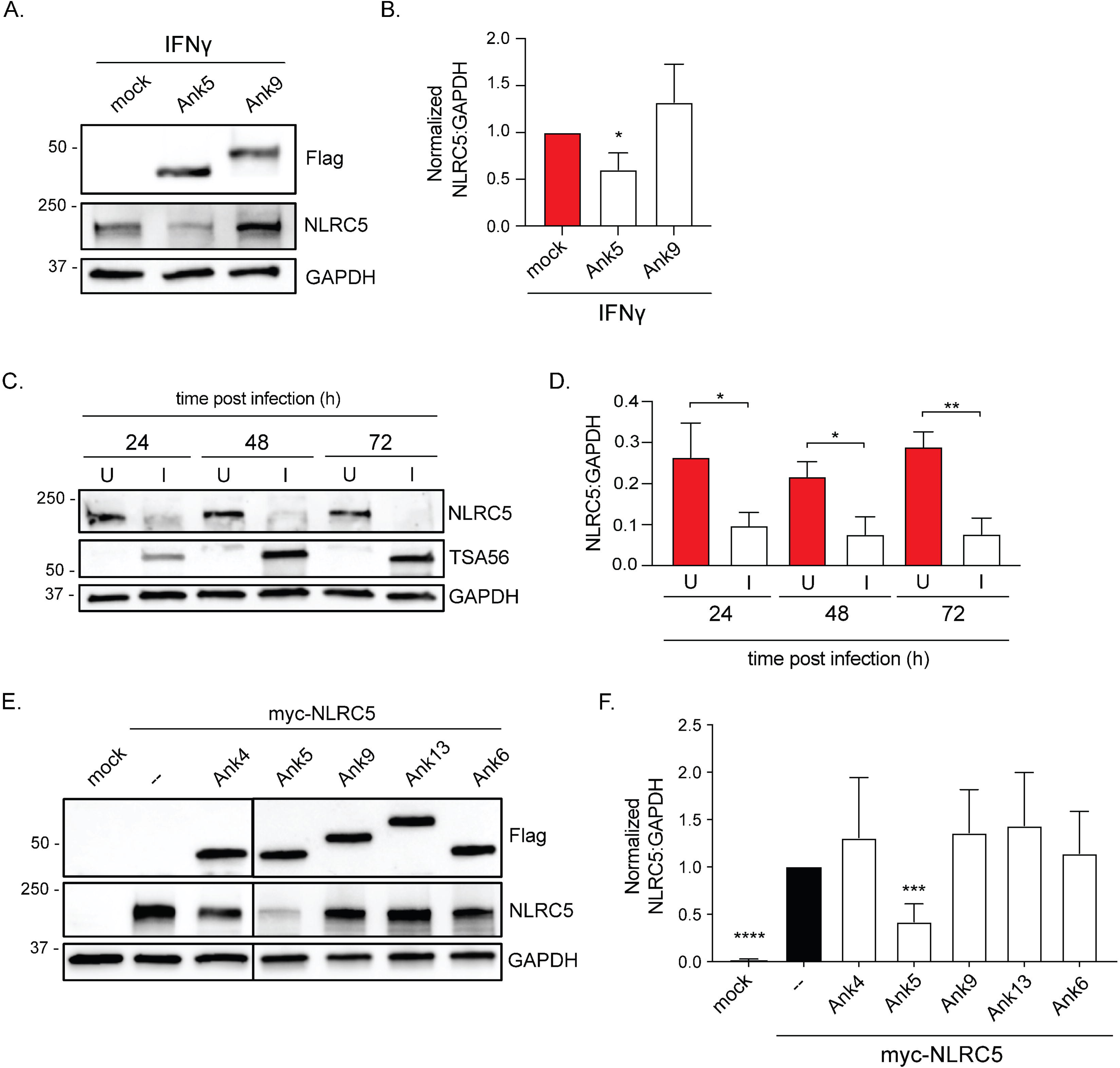
Ectopically expressed Ank5 and *O. tsutsugamushi* promote NLRC5 degradation. (A and B) EA.hy926 cells were mock transfected or transfected to express Flag-Ank5 or Flag-Ank9 and treated with IFNγ. (A) Whole cell lysates were analyzed by immunoblotting. (B) Densitometric quantification of the NLRC5:GAPDH signal ratios from input blots. Data are means ± SD (n = 3). An unpaired *t-*test was used to determine statistical significance among pairs compared to mock transfected cell (red bar). (C and D) EA.hy926 cells were mock [U] or infected with *O. tsutsugamushi* [I] followed by IFNγ treatment. (C) Whole cell lysates were analyzed by immunoblotting. (D) Densitometric quantification of the NLRC5:GAPDH signal ratios. Data are means ± SD (n = 3). An unpaired *t*-test determined statistical significance among uninfected and infected pairs. (E and F) HeLa cells were mock transfected or transfected to express myc-NLRC5 alone (--) or together with Flag-tagged Ank4, Ank5, Ank9, Ank13, or Ank6. (E) Whole cell lysates were analyzed by immunoblotting. Vertical lines between bands indicate where the blot was cropped or imaged separately. (F) The densitometric signal value of NLRC5 was divided by the corresponding GAPDH densitometric signal value and normalized to the densitometric signal level of myc-NLRC5 expressing cells (--; black bar). Data are means ± SD (n = 4). An unpaired *t-*test was used to determine statistical significance among pairs compared to myc-NLRC5 (--; black bar). *, p < 0.05; **, p < 0.01; ***, p < 0.001; ****, p < 0.0001.

**Extended Data Figure 5.**
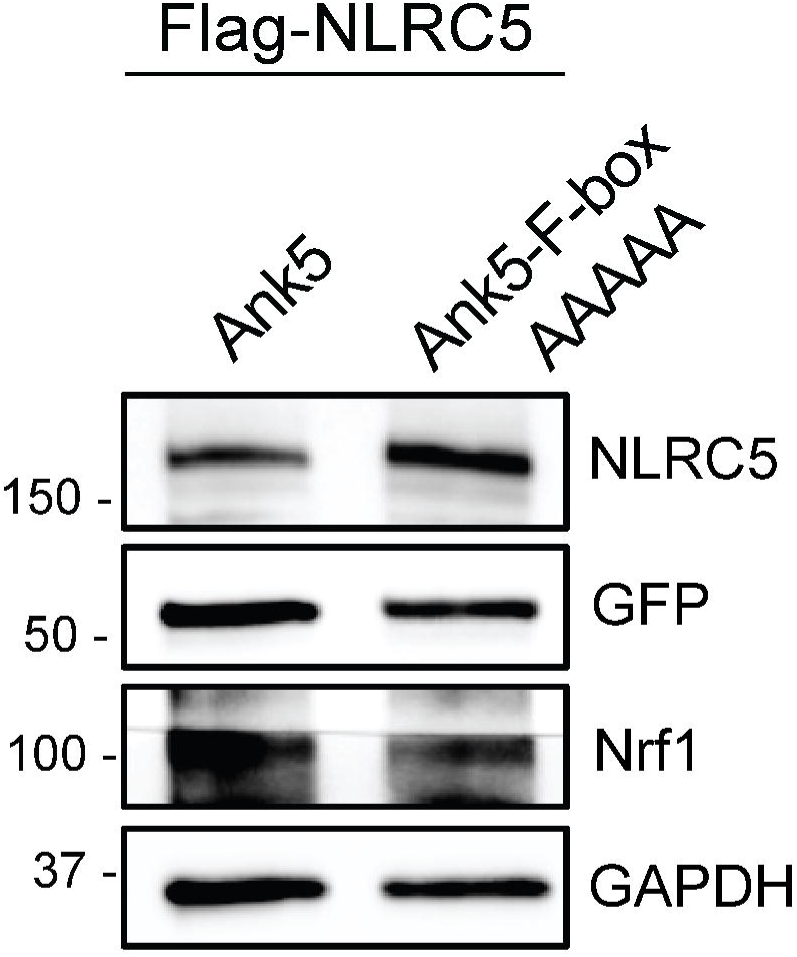
Input lysates from cells expressing GFP-tagged Ank5 and Ank5-F-boxAAAAA expressing cells for LC-MS/MS IP-Proteomics. Western blots of input lysates for LC-MS/MS IP-Proteomics to verify successful MG132 treatment and expression of all proteins of interest by cells that were transfected to co-express Flag-NLRC5 and either GFP-Ank5 or GFP-Ank5-F-boxAAAAA.

**Extended Data Figure 6.**
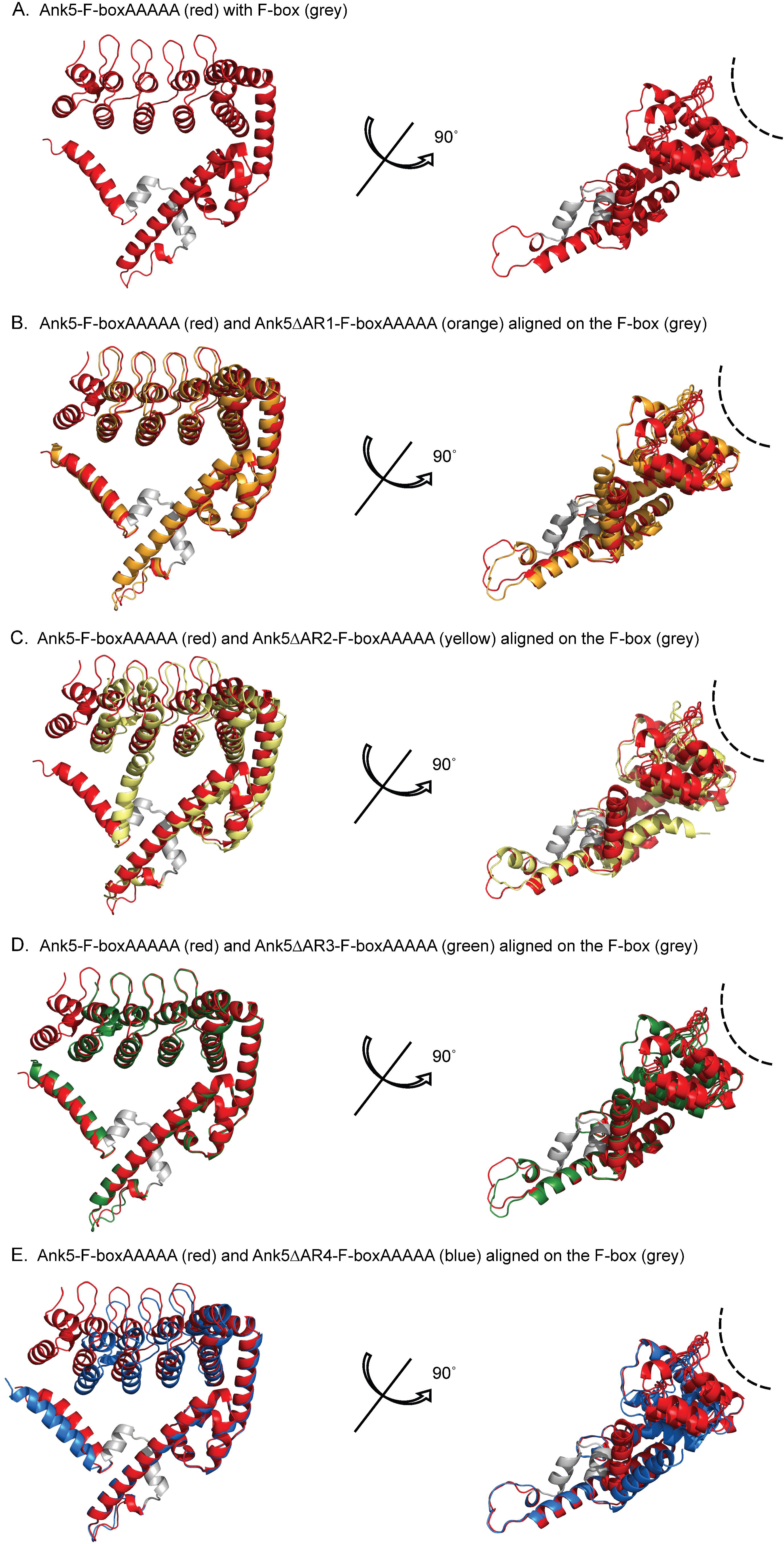
Predicted structures for Ank5-F-boxAAAAA and ΔAR mutants thereof. (A–E) AlphaFold was used to predict structures for Ank5-F-boxAAAAA, Ank5-ΔAR1-F-boxAAAAA, Ank5-ΔAR2-F-boxAAAAA, Ank5-ΔAR3-F-boxAAAAA, and Ank5-ΔAR4-F-boxAAAAA. (A) Predicted structure for Ank5-F-boxAAAAA. (B to D) Predicted structures for Ank5-ΔAR1-F-boxAAAAA (B), Ank5-ΔAR2-F-boxAAAAA (C), Ank5-ΔAR3-F-boxAAAAA (D), and Ank5-ΔAR4-F-boxAAAAA (E) were overlaid on Ank5-F-boxAAAAA. For each overlay, the structures were aligned to the F-box of Ank5-F-boxAAAAA (colored gray). The interacting groove formed by the AR domain is indicated by the dashed line arc.

**Extended Data Figure 7.**
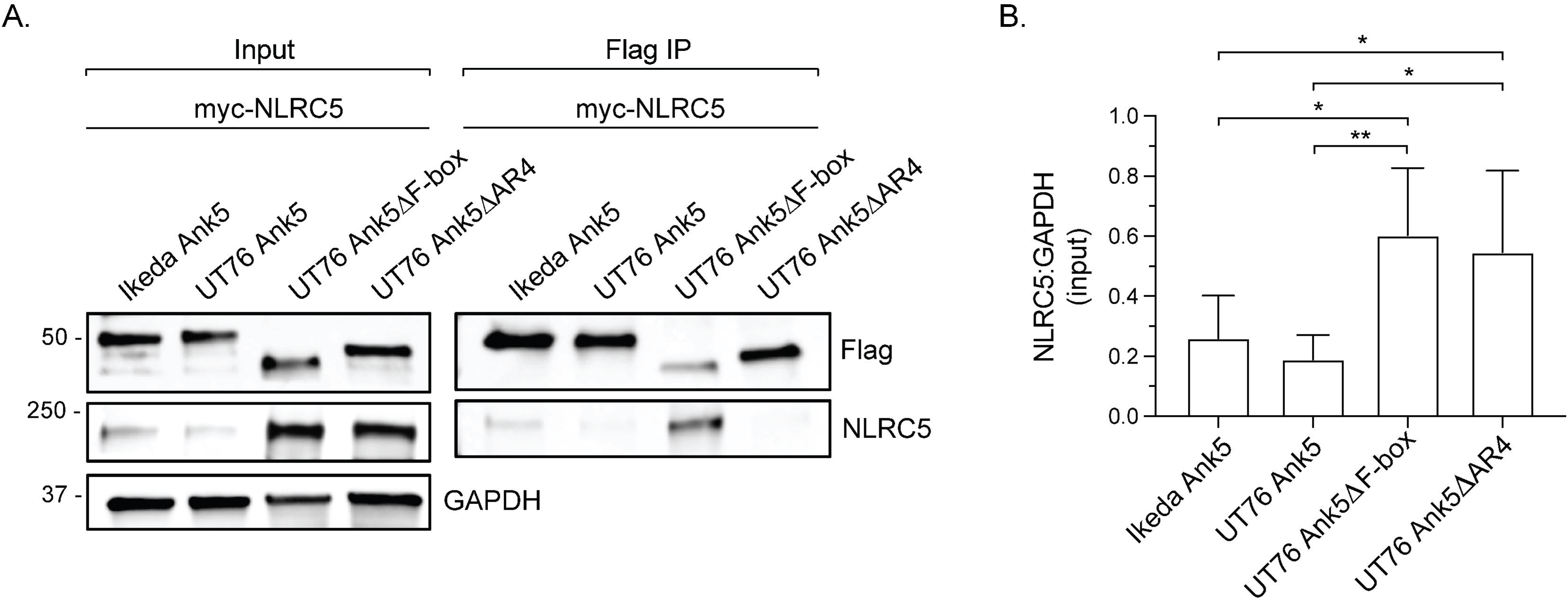
UT76 Ank5 AR4 binds NLRC5 and directs NLRC5 degradation using its F-box. (A and B) HeLa cells were transfected to co-express Flag-Ikeda Ank5, -UT76 Ank5, -UT76 Ank5ΔF-box, or -UT76 Ank5ΔAR4 with myc-NLRC5. (A) Input lysates and immunoprecipitated Flag-tag protein (Flag IP) complexes were analyzed by immunoblotting. (B) Densitometric quantification of the NLRC5:GAPDH signal ratio from input blots. Data are means ± SD (n = 7). One-way ANOVA with Tukey’s *post hoc* test was used to assess significant differences between every mean. *, p < 0.05; **, p < 0.01.

**Extended Data Figure 8.**
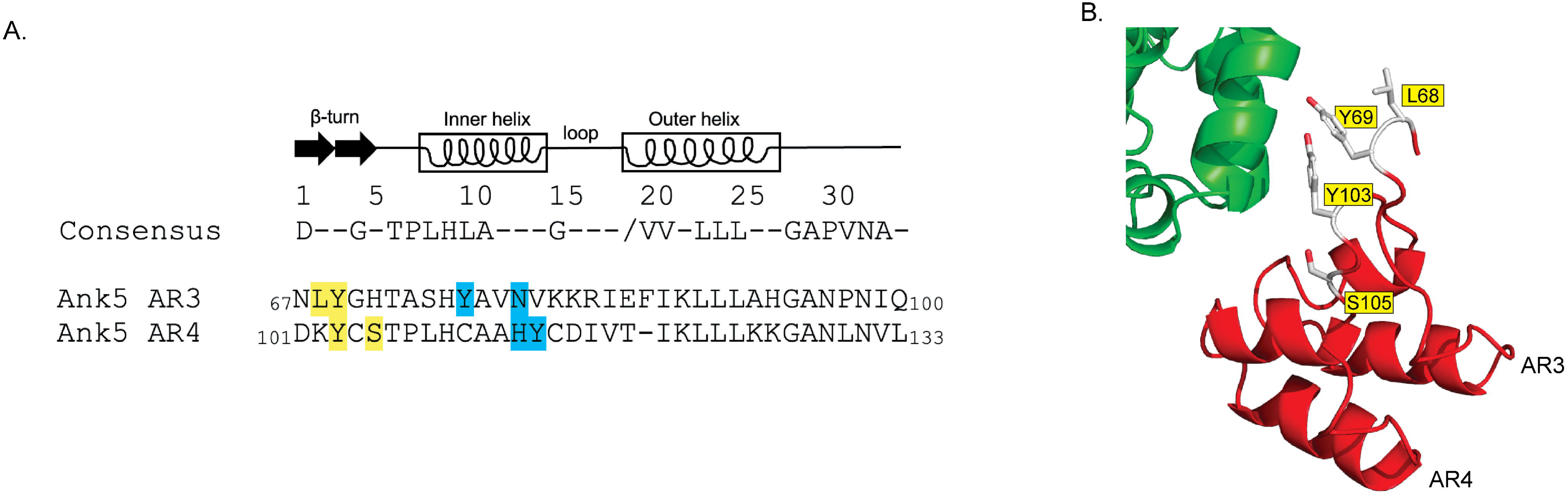
In silico prediction of Ank5 residues that interact with NLRC5. (A) NovaDock predicted that Ank5 residues in AR3 and AR4 bind NLRC5. Presented are the amino acid sequences of AR3 and AR4 aligned with the consensus sequence for a eukaryotic AR and a schematic for AR secondary structure. Residues highlighted yellow (AR3 L68 and Y69; AR4 Y103 and S105) occur at positions 2, 3, and/or 5 in the β-turn that tend to mediate protein-protein interactions. Those highlighted cyan are at positions in the inner α-helix that would alter AR tertiary structure if mutated. (B) Novadock modeled interaction between Ank5 and NLRC5. The secondary structures of Ank5 AR3 and AR4 are red while that of the NLRC5 NACHT domain is green. The Ank5 residues predicted to interact with NLRC5 and that were targeted for alanine substitution are labeled.

**Extended Data Table 1.** NLRC5 interactome recovered from *O. tsutsugamushi* infected cells*^a^*.

| Proteins recovered | GO term<sup>b</sup> (node color<sup>c</sup>) | Protein count |
| --- | --- | --- |
| LGALS1, HNRNPL, RPL18A, C1QBP, DDX5, YBX3, RPS12, HSPE1, DDX39A, CANX, RPS16, TRIM28, C14orf166, SRP14, CCT6A, NONO, CCT5, HNRNPU, KPNB1, CCT3, RPN1, PPIB, PDIA3, RPS9, FASN, SRP9, RPL13, EEF2, RPL38, RPL4, PABPC1, TCP1, NCL, RPS15A, IMPDH2, PLEC, PRPF31, P4HB, RBM10, RPS27L, DHX15, GANAB, RPL27A, RPL21, RPS17, DYNC1H1, SPTBN1, STIP1, XRCC6, EZR, SERBP1, HNRNPH2, RPS4X, RPL10A, ANXA1, RBBP7, RPS6, DUT, RPL13A, ACTN1, HLA-A, RPS8, DDX3X, PCBP3, TARS, STRAP, RPL17, RPL6, EEF1D, RPS24, PSMA1, FLNB, GNB2L1, RPL30, RPS13, RAN, RPL18, RPL27, CLTC | RNA binding (yellow) | 79 |
| RPL18A, RPS12, RPS16, RPS9, RPL13, EEF2, RPL38, RPL4, PABPC1, RPS15A, RPS27L, RPL27A, RPL21, RPS17, WARS, RPS4X, RPL10A, RPS6, RPL13A, RPS8, DDX3X, TARS, RPL17, RPL6, EEF1D, RPS24, GNB2L1, RPL30, RPS13, RPL18, RPL27 | Translation (blue) | 31 |
| RPL18A, RPS12, SSR1, RPS16, SRP14, CALM3, RPS9, SRP9, RPL13, RPL38, RPL4, RPS15A, RPL27A, RPL21, RPS17, SPTBN1, RPS4X, RPL10A, BAG6, RPS6, RPL13A, RPS8, RPL17, RPL6, RPS24, RPL30, RPS13, RPL18, RPL27 | Establishment of protein localization to membrane (red) | 29 |
| CNN2, TMPO, FASN, EEF2, RUVBL1, NUDC, PLEC, P4HB, BSG, SFN, PRDX6, SPTBN1, EZR, BAG3, SERBP1, SPTAN1, CTTN, ANXA1, ACTN1, DDX3X, IGF2, CAPZB, RPL6, EEF1D, FLNB, GNB2L1, RAN | Cell adhesion molecule binding (magenta) | 27 |
| CNN2, TMPO, FASN, EEF2, RUVBL1, NUDC, PLEC, BSG, SFN, PRDX6, SPTBN1, EZR, BAG3, SERBP1, SPTAN1, CTTN, ANXA1, DDX3X, CAPZB, RPL6, EEF1D, FLNB, GNB2L1, RAN | Cadherin binding (light green) | 24 |
| RPL18A, RPS12, RPS16, RPS9, RPL13, RPL38, RPL4, RPS15A, RPS27L, RPL27A, RPL21, RPS17, RPS4X, RPL10A, RPS6, RPL13A, RPS8, RPL17, RPL6, RPS24, RPL30, RPS13, RPL18, RPL27 | Ribosome (cyan) | 24 |
| HSPE1, CANX, CCT7, CCT6A, AIP, CCT5, CCT3, CCT2, PPIB, CLU, TCP1, NUDC, CCT6B, DNAJB11, RUVBL2 | Unfolded protein binding (orange) | 15 |
| PSMA4, PSMD3, PSMA5, PSMD1, PSMD2, PSMC5, PSMA7, PSME1, PSMC2, PSMA1 | Proteasome (dark green) | 10 |

<sup>a</sup>Flag-NLRC5 interacting proteins from cells infected with *O. tsutsugamushi* relative to uninfected cells
<sup>b</sup>GO, gene ontology
<sup>c</sup>As presented in Figure S2.

**Extended Data Table 2.**
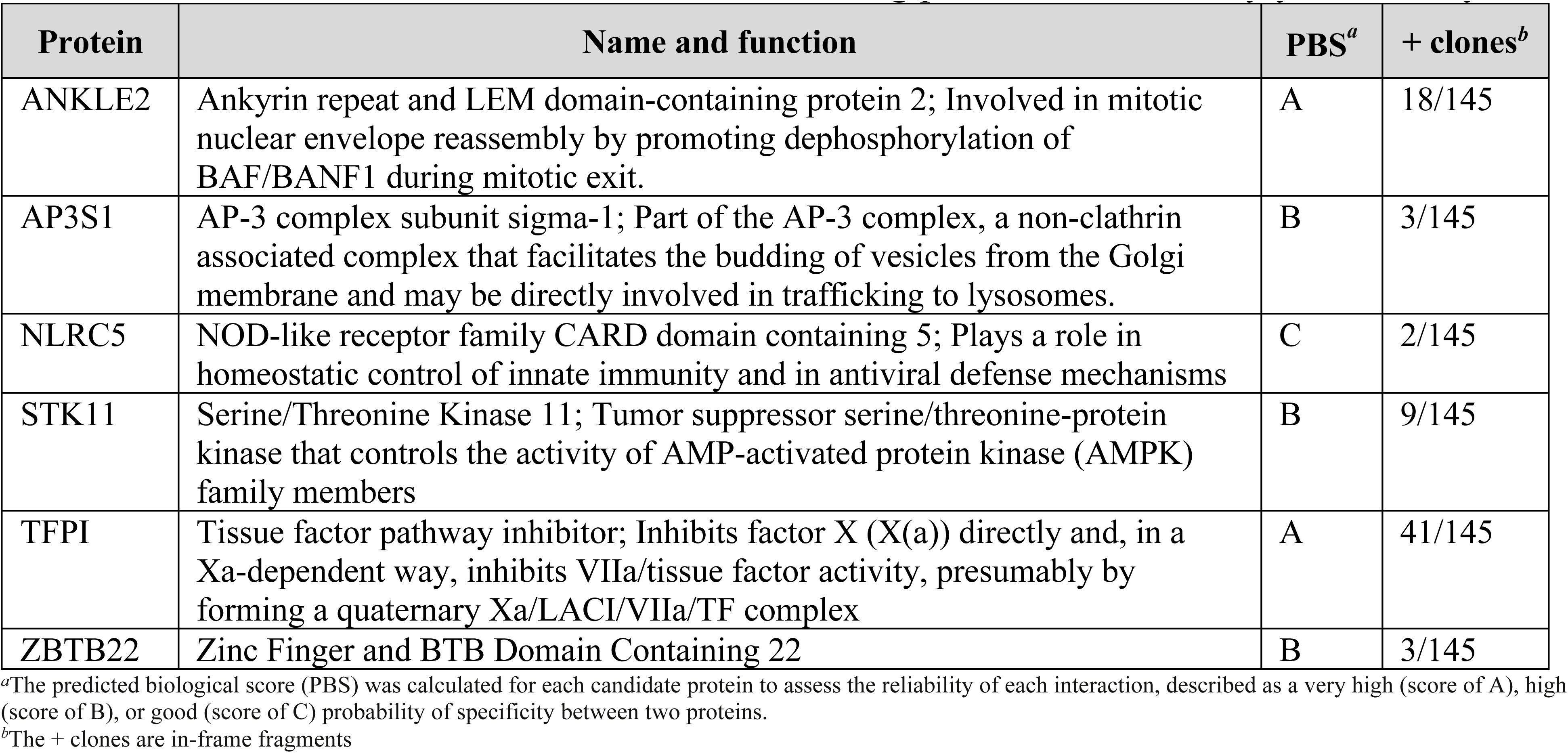
Candidate Ank5ΔF-box binding partners identified by yeast two-hybrid.

| Protein | Name and function | PBS<sup>a</sup> | + clones<sup>b</sup> |
| --- | --- | --- | --- |
| ANKLE2 | Ankyrin repeat and LEM domain-containing protein 2; Involved in mitotic nuclear envelope reassembly by promoting dephosphorylation of BAF/BANF1 during mitotic exit. | A | 18/145 |
| AP3S1 | AP-3 complex subunit sigma-1; Part of the AP-3 complex, a non-clathrin associated complex that facilitates the budding of vesicles from the Golgi membrane and may be directly involved in trafficking to lysosomes. | B | 3/145 |
| NLRC5 | NOD-like receptor family CARD domain containing 5; Plays a role in homeostatic control of innate immunity and in antiviral defense mechanisms | C | 2/145 |
| STK11 | Serine/Threonine Kinase 11; Tumor suppressor serine/threonine-protein kinase that controls the activity of AMP-activated protein kinase (AMPK) family members | B | 9/145 |
| TFPI | Tissue factor pathway inhibitor; Inhibits factor X (X(a)) directly and, in a Xa-dependent way, inhibits VIIa/tissue factor activity, presumably by forming a quaternary Xa/LACI/VIIa/TF complex | A | 41/145 |
| ZBTB22 | Zinc Finger and BTB Domain Containing 22 | B | 3/145 |

<sup>a</sup>The predicted biological score (PBS) was calculated for each candidate protein to assess the reliability of each interaction, described as a very high (score of A), high (score of B), or good (score of C) probability of specificity between two proteins.
<sup>b</sup>The + clones are in-frame fragments

**Extended Data Table 3.** Comparison of full-length Ank5 homologs in other *O. tsutsugamushi* strains/isolates.

| Strain/<br>isolate | Geographic<br>Origin | Reference | Nucleotide<br>length | %<br>Nucleotide<br>Identity | Locus Tag | AA<br>Length | % AA<br>Identity | Number<br>of ARs | % AA<br>Identity<br>AR4 |
| --- | --- | --- | --- | --- | --- | --- | --- | --- | --- |
| Ikeda | Japan | Tamura 1984 | 1038 | 100 | OTT_RS01000<sup>a</sup> | 346 | 100 | 4 | 100 |
| Ikeda | Japan | Tamura 1984 | 1038 | 100 | OTT_RS01950 | 346 | 100 | 4 | 100 |
| Ikeda | Japan | Tamura 1984 | 1038 | 100 | OTT_RS02325 | 346 | 100 | 4 | 100 |
| UT76 | NE Thailand | Blacksell<br>2008 | 1041 | 96 | UT76HP_01926 | 347 | 92 | 4 | 100 |
| Wuj/2014 | China |  | 915 | 97 | CP044031 | 304 | 90 | 4 | 100 |

<sup>a</sup>Copy selected to represent all three Ank5 copies and to which homologs were compared (VieBrock et al., 2015).

