## Supplementary Table 1 for "*Orientia tsutsugamushi* Ank5 directs ubiquitination and proteasomal degradation of NLRC5 to inhibit major histocompatibility complex class I expression"

Supplementary Table 1. Oligonucleotides utilized in this study

| **Designation**[a] | **Sequence (5'-3')** |
|---|---|
| *tsa56*_715F | GTTACTGCATTGTCACATGCTAAT |
| *tsa56*_835R | GAACTTCAATATTAGCTACCTTAGCGA |
| *tsa56*_Full_F | CTTTTAAGGAGATTAGAATGAAAAAAATTATGT |
| *tsa56*_Full_R | AGAAAAACTAGAAGTTATAGCGTACAC |
| *ank5*-318F | TCCTCTTCATTGCGCTGCTCATTAC |
| *ank5*-539R | TCACTTCCACTAATCTTTATGCCAAGATGCTC |
| *ott16S*-911F | GTGGAGCATGCGGTTTAATTCGATGATC |
| *ott16S*-1096R | TAAGAATAAGGGTTGCGCTCGTTGC |
| Ank5ΔAR1 F[b] | *CA*ATGATGGACATGACCGGCACCACC |
| Ank5ΔAR1 R | TCATGTCCATCATTGAATTCGCGGCCG |
| Ank5ΔAR2 F | ACCTGGGTAATTTGTACGGCCACACC |
| Ank5ΔAR2 R | ACAAATTACCCAGGTTGATGTAGTCC |
| Ank5ΔAR3 F | ACATCCAAGACAAGTACTGTTCCACCCC |
| Ank5ΔAR3 R | ACTTGTCTTGGATGTTCGGGTTGGC |
| Ank5ΔAR4 F | ACATTCAGGACGTGACCCACAGCACC |
| Ank5ΔAR4 R | TCACGTCCTGAATGTTGGGATTGGCGCC |
| Ank5$_{L68A}$ F[c] | CCAAAATGCCTACGGCCACACCGCAAGC |
| Ank5$_{L68A}$ R[c] | CCGTAGGCATTTTGGATGTTCGGGTTGGCTCC |
| Ank5$_{Y69A}$ F[c] | AAATTTGGCCGGCCACACCGCAAGCCAT |
| Ank5$_{Y69A}$ R[c] | TGGCCGGCCAAATTTTGGATGTTCGGGTTGGCT |
| Ank5$_{Y103A}$ F[c] | GGACAAGGCCTGTTCCACCCCACTCCATTG |
| Ank5$_{Y103A}$ R[c] | GAACAGGCCTTGTCCTGAATGTTGGGATTGGC |
| Ank5$_{S105A}$ F[c] | GTACTGTGCCACCCCACTCCATTGCGCA |
| Ank5$_{S105A}$ R[c] | GGGGTGGCACAGTACTTGTCCTGAATGTTGGGA |

[a]F and R refer to primers that bind to the sense and antisense strand, respectively.
[b]Italicized text indicate nucleotides complementary to the p3XFLAG-CMV-7.1 vector.
[c]Highlighed text indicate nucleotides used to make site-directed mutants.
